# Facilitating taxonomy and phylogenetics: An informative and cost-effective protocol integrating long amplicon PCRs and third generation sequencing

**DOI:** 10.1101/2023.08.03.551825

**Authors:** Domagoj Gajski, Jonas O. Wolff, Anja Melcher, Sven Weber, Stefan Prost, Henrik Krehenwinkel, Susan R. Kennedy

**Affiliations:** Department of Biogeography, Faculty of Spatial and Environmental Sciences, University of Trier, Universitätsring 15, Trier 54296, Germany; Department of Botany and Zoology, Faculty of Science, Masaryk University, Kotlářská 2, Brno 611 37, Czech Republic; Evolutionary Biomechanics, Zoological Institute and Museum, University of Greifswald, Loitzer Str. 26, Greifswald 17489, Germany; School of Natural Sciences, Macquarie University, NSW 2109, Sydney, Australia; Ecology and Genetics Research Unit, University of Oulu, Pentti Kaiteran katu 1, Linnanmaa, Finland

**Keywords:** spider, taxonomy, phylogeny, nanopore sequencing, PCR primers, DNA barcoding

## Abstract

Phylogenetic inference has become a standard technique in integrative taxonomy and systematics, as well as in biogeography and ecology. DNA barcodes are often used for phylogenetic inference, despite being strongly limited due to their low number of informative sites. Also, because current DNA barcodes are based on a fraction of a single, fast-evolving gene, they are highly unsuitable for resolving deeper phylogenetic relationships due to saturation. In recent years, methods that analyse hundreds and thousands of loci at once have improved the resolution of the Tree of Life, but these methods require resources, experience and molecular laboratories that most taxonomists do not have. This paper introduces a PCR-based protocol that produces long amplicons of both slow- and fast-evolving unlinked mitochondrial and nuclear gene regions, which can be sequenced by the affordable and portable ONT MinION platform with low infrastructure or funding requirements. As a proof of concept, we inferred a phylogeny of a sample of 63 spider species from 20 families using our proposed protocol. The results were overall consistent with the results from approaches based on hundreds and thousands of loci, while requiring just a fraction of the cost and labour of such approaches, making our protocol accessible to taxonomists worldwide.

**Highlights:**

- DNA barcoding is an invaluable tool for fast and accurate taxonomic classification
- Existing DNA barcodes are still insufficient for obtaining well-supported phylogenies
- We present a protocol that produces long amplicons of unlinked loci for spiders
- Amplicons are sequenced at very low cost per specimen with ONT MinION
- Our recovered phylogeny is largely consistent with that of high-cost approaches.

## 1. Introduction

In the past few decades, DNA barcoding has emerged as an invaluable tool for the fast and accurate identification of taxa (Antil et al. 2023; Jones et al., 2021; Banchi et al. 2020; Ratnasingham & Hebert, 2007). Provided that an adequate reference database is available, DNA barcodes can be used to reliably identify organisms, and even facilitate taxonomic discovery by detecting as-yet uncharacterized taxa (Valentini et al., 2009; Hamilton et al., 2011). DNA barcodes can serve as an important complement to traditional taxonomy (Chan et al. 2014), assisting, for example, with identifying taxa that are challenging due to high levels of trait variation (Lopez-Vaamonde et al. 2021). For hyperdiverse taxa, such as arthropods, DNA barcoding has become indispensable because traditional morphological identification often poses an insurmountable challenge in terms of expertise and time required (Hajibabaei et al., 2016). At the same time, accurately identifying such taxa is critical for ecological and conservation research (Costello et al., 2015; Engel et al., 2021), making DNA barcoding highly important.

Besides their advantages for taxonomic identification, DNA barcodes can also provide phylogenetic information and are often used in integrative taxonomy to identify monophyletic groups and classify genera and species. Knowledge of evolutionary relationships can offer deeper insights into biological communities, potentially allowing inferences about functional traits or ecological relationships (Kress et al., 2015; Xie et al., 2023). With the advent of high-throughput sequencing (HTS), the value of PCR-based DNA barcoding has increased even further. Barcodes can now be generated for thousands of specimens in parallel at a fraction of the labor and cost previously needed (Shokralla et al., 2015; Slatko et al., 2018). Numerous DNA barcoding markers have been established across the tree of life (Hebert et al., 2003; Schoch et al., 2012), with some having large sequence reference databases (e.g. COI barcode in Barcode of Life Data System; Ratnasingham & Hebert, 2007).

Despite the substantial benefits of DNA barcoding, the methodology still has limitations. DNA barcodes - especially uniparentally inherited markers such as those located in the mitochondrial compartment - can lead to over- or under-spliting splitting when species have high levels of male-biased gene flow (Krehenwinkel et al., 2016), infections with reproductive parasites (e.g. *Wolbachi*a and *Cardinium*; Hurst & Jiggins, 2005) or introgressive hybridization between one another (Melo-Ferreira et al., 2005). Furthermore, DNA barcodes are usually relatively short, up to several hundred bp (Tan et al., 2019; Shendure & Ji, 2008). This limits their number of potential phylogenetically informative sites, which in turn decreases their utility for understanding evolutionary relationships. These relationships, however, are the very foundation of the taxonomic classification system (Hinchliff et al. 2015). In some cases, short DNA barcodes do not provide enough information to distinguish species pairs with relatively recent divergence (Meier et al., 2006; Stallman et al., 2019). Independent of barcode length, another fundamental issue is that the evolutionary history of a single gene might not mirror that of the entire genome, let alone the entire species (Rubinoff & Holland, 2005; Winkler et al., 2015). For instance, fast-evolving genes will reach saturation before the divergence times of older clades, making them unsuitable for the analysis of any but relatively shallow evolutionary divergences (Arbogast et al., 2002). Thus, an ideal barcode or set of barcodes should include a combination of fast- and slow-evolving genes, ideally with at least some nuclear (biparentally inherited) loci (Funk & Omland, 2003).

In recent years, many studies have circumvented these problems by analysing whole transcriptome sequences (Wong et al., 2019), whole genomes (Zhang et al., 2016), ultraconserved elements (UCE) by targeted enrichment (Faircloth et al., 2012), or thousands of anonymous loci obtained by ddRAD sequencing (Ortiz et al., 2021). These advances have significantly improved the resolution of the tree of life, even within deeply divergent and challenging clades (Kulkarni et al., 2020; Misof et al., 2014). Yet such high-throughput methods have two main problems: the high cost of generating data and the need for a well-equipped molecular laboratory. The majority of the world’s taxonomists lack direct experience, resources or access to molecular laboratories that are set up for genome-level taxonomic analysis.

Given this context, a DNA barcoding approach would be preferable which increases the length of barcode sequences (and the number of informative sites), targets several unlinked loci across the genome, and can still remain reasonably straightforward and affordable to taxonomists all around the world. A highly promising solution for the affordability presents itself with Nanopore sequencing (Oxford Nanopore Technologies (ONT)), which provides inexpensive, portable and straightforward third-generation sequencing (Wang et al., 2021). These platforms are perfectly suited for targeting long-read-length PCR amplicons and can process hundreds of samples in a single sequencing run. Despite a relatively high raw read error rate (up to 10%; Delahaye & Nicolas, 2021), one can derive highly accurate consensus sequences of the studied amplicons at a relatively low sequencing depth (Pomerantz et al., 2018, Krehenwinkel et al.2019).

Here, we created a new barcoding approach using a newly designed set of PCR primers and sequencing on ONT’s MinION platform to obtain long amplicons of mitochondrial and nuclear gene regions across the spider tree of life. We focused on spiders due to their extremely high diversity (over 50,000 species described globally (World Spider Catalog, 2023) and many more undescribed (Agnarsson et al. 2013)) and the fact that their morphological identification is notoriously challenging, especially when diagnostic features - typically the genitalia - are damaged or missing (e.g. in juvenile specimens; Kuntner, 2022). Although here we optimized the primers for spiders as our model system with a few nucleotide adjustments, this protocol could be implemented for many other arthropod clades, making our approach widely applicable. To test the efficacy of our protocol, we performed a case study with spider specimens collected in a semi-arid grassland in western Germany and at several locations in eastern Australia and the South Island of New Zealand. The 129 collected specimens represent 63 species in 47 genera and 20 families, covering major evolutionary lineages across the spider tree of life (Synspermiata, Araneoidea, Marronoid Clade, Oval Calamistrum Clade, and Dionycha). We tested the efficacy of our protocol for obtaining accurate barcodes from this diverse assemblage of spiders. Additionally, we tested the extent to which our markers could recover a phylogeny for these 63 species that agrees with current phylogenetic hypotheses built from approaches that either used several short amplicons (Wheeler et al. 2017) or hundreds of thousands of loci (Azevedo et al., 2022; Fernández et al., 2018; Kulkarni et al., 2021; 2022). Lastly, we tested whether our long amplicons are more or less effective than the standard, 658-bp COI barcode (Barrett & Hebert, 2005) at identifying a barcode gap - i.e., a specified range of phylogenetic distances between species within a genus.

## 2. Methods

### 2.1 Design of DNA-barcoding primers

For the amplification of mitochondrial genes, we focused on Cytochrome C Oxidase I (COI) and Cytochrome B (CytB), two genes commonly used for barcoding and phylogenetics of spiders and other metazoans (Parson et al., 2000; Wheeler et al., 2017). Our aim was to design primer pairs to amplify large fragments (>= 700 bp) in order to maximize information yield. The maximal potential amplicon size for design was limited by the natural size of the gene.

Using the NCBI Nucleotide database (accessed May 2021; Sayers et al., 2022), we searched for all mitochondrial genomes belonging to spiders. The following search terms were used: (((mitochondrial[All Fields] AND genome[All Fields]) AND “animals”[porgn]) AND “arthropods”[porgn]) AND “spiders”[porgn] AND (“10000”[SLEN] : “20000”[SLEN]). This way, we found only mitochondrial genomes belonging to spiders with the appropriate genome size between 10,000 and 20,000 bp, a common size range of mitochondrial genomes for all animals (Boore, 1999). For each available result, we downloaded the COI and CytB sequences as fasta files.

For each respective gene, the downloaded sequences were aligned using the Clustal Omega alignment tool (Sievers et al., 2011) in Geneious 2020.0.02. We then visually identified conserved regions inside the aligned sequences where we could design new primers. We selected primer sites based on the following criteria: min. size of 18 bp, min. melting temperature (Tm) of 45 °C, GC-content between 20 and 80 %, number of degenerate bases no more than 6 (i.e. no more than once every three bases), and at least one of the first three bases on the 3’ site of the primer should be either G or C (to increase the binding strength of the primer). We focused mainly on the beginning and end of a gene’s sequence to maximize the length of the resulting amplicon. Lastly, combinations of forward and reverse primers were made based on the similarity of the Tm, with the difference never exceeding 5 °C.

For nuclear barcodes, we focused on the ribosomal cluster, which spans from the 18S rRNA-encoding gene (18S) to the 28S rRNA-encoding gene (28S). This region also includes the internal transcribed spacers (ITS1 and ITS2) and the 5.8S rRNA gene. This ensured the inclusion of both highly conserved regions (18S, 5.8S and parts of 28S) and the fast-evolving internal transcribed spacers (ITS1 and ITS2), providing informativeness at both shallow and deeper phylogenetic levels (Krehenwinkel et al., 2019). Additionally, this region is generally known for its efficiency in barcoding and phylogenetic research, and a recent study sequenced both 18S and 28S for a diverse range of spiders representing most of the spider tree of life (Wheeler et al., 2017). Thus, rather than searching all of NCBI, we used 18S and 28S sequences of spiders found in Wheeler et al. (2017). Sequences were aligned, and primer sites were identified as described above. Here we designed forward primers only in 18S and reverse primers in 28S, thereby targeting an amplicon spanning the whole ribosomal cluster. Combinations of forward and reverse primer were made based on the similarity of the Tm, with the difference never exceeding 5 °C. Lastly, we compared the efficiency of these primers with the primer pair 18S_F4 and 28S_R8 for the 18S-28S amplicon already developed for a previous study (Krehenwinkel et al., 2019.)

All primers were designed with unique 20-bp index sequences attached to the 5’- end to allow for demultiplexing by sample after sequencing. These indexes were designed using Barcode Generator (http://comailab.genomecenter.ucdavis.edu/index.php/Barcode_generator, accessed August 2021) with a minimum distance of 10 bases between each index (as shown in Pomerantz et al. 2022).

### 2.2 Optimization of the PCR protocol

An exhaustive series of PCR trials was performed on 25 adult spider specimens belonging to 24 distinct species and 15 different families (Tab. A.1) to identify the best-performing primers for each gene. Specimens were collected by hand on a semi-arid grassland at Hohengöbel near Kimmlingen, Germany (49°49’58.5”N 6°36’05.6”E; permit # 425-104-235-005/2021 from the Struktur- und Genehmigungsdirektion Nord of Rheinland-Palatinate) and stored in individual tubes filled with absolute ethanol. Spider tissues were macerated with a Mini-Beadbeater 16 (BioSpec Products, Bartlesville, OK, USA), using 3-mm stainless steel ball bearings (Viwanda), for 1-2 minutes or until the spiders were thoroughly pulverized. DNA was then extracted using the Qiagen PureGene tissue kit following the manufacturer’s instructions, with an overnight (16-20 h) cell lysis at 55 °C. PCR trials were run on the DNA extracts using the Qiagen Multiplex PCR kit (Qiagen, Hilden, Germany), following the manufacturer’s protocol. In addition to identifying the most effective primer pairs, we assessed the optimal annealing temperature (using gradient PCR), the optimal primer concentration, and whether multiple markers could be amplified together in a multiplex PCR.

### 2.3 Case study

To test the efficacy of our protocol for recovering evolutionary relationships among spiders, we collected an additional 104 specimens to add to our existing dataset of the 25 individuals used for PCR optimization. Specimens were collected at the same location as above by hand, branch beating, or net sweeping, and immediately stored individually in absolute ethanol. Specimens were afterwards identified to species level using identification keys (Nentwig et al., 2022). Additionally, we included specimens from nine spider species (representing three families) endemic to Australia and New Zealand that had been collected in 2019 (under permits SL101868, FA18285, 71225-RES) and stored in absolute ethanol at -20° C. When possible, several specimens were included per species to check for intraspecific differences in our barcoding markers. Our final data set consisted of 129 specimens belonging to 63 species and 47 genera in 20 families (Tab. A.2). Tissue was macerated, and DNA was extracted as described above.

Based on the results of the PCR optimization tests, we chose primer sets that amplify large fragments of the mitochondrial genes COI and CytB, as well as the nuclear 18S-28S region, across the spider tree of life (Tab. 1). The applicability of these primers across a broad range of spider taxa is highlighted in Fig. 1. To reduce cost and effort, COI and CytB were amplified together in a multiplex PCR, using the Qiagen Multiplex kit following the manufacturer’s protocol, with 30 cycles and an annealing temperature of 48 °C. The CytB amplicon is, on average, 358 bp shorter than the COI amplicon (696 vs. 1054 bp) and was consequently expected to amplify preferentially. For that reason, and based on the results of our PCR trials, we doubled the concentration of COI primers relative to CytB primers in the multiplex. The 10-µL volume of each reaction thus consisted of 5 µL of Qiagen Multiplex mix, 0.5 µL each of 10-µM CytB primers, 1 µL each of 10-µM COI primers, 1 µL of PCR water, and 1 µL of template DNA. In addition, we amplified a ∼4,500 bp fragment of the nuclear ribosomal cluster (from the 18S rRNA-encoding to the 28S rRNA-encoding regions) using primers 18S_F4 and 28S_R8 published in Krehenwinkel et al. (2019; Fig. 1). The length of this amplicon (∼4.5 kb) was expected to pose a challenge to the Qiagen Multiplex kit, which is optimized for fragments up to 1.5 kb (Qiagen Multiplex kit handbook 10/2010). Thus, to improve the yield of the 18S-28S amplicon, we used the Qiagen UltraRun Long-Range kit and followed the manufacturer’s recommendations for cycling conditions (3 minutes initial denaturation at 93 °C, followed by 40 cycles of 30 s denaturation at 93 °C and 2 min annealing/extension at 68 °C, with a final extension step of 10 min at 72 °C). Successful amplification was verified by electrophoresis on 1 % agarose gels.

**Figure 1.**
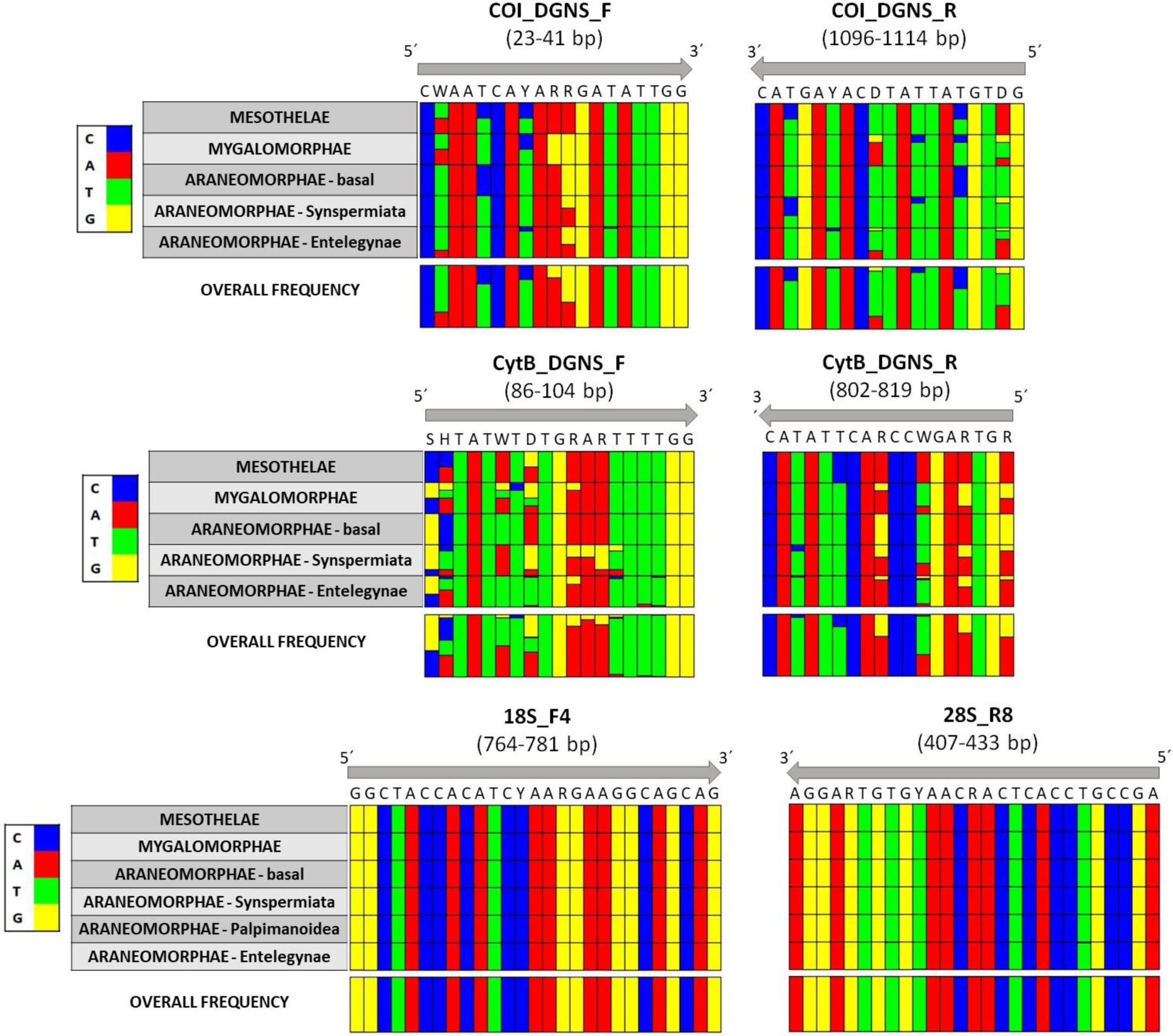
The nucleic acid composition of every base position inside the COI_DGNS_F and COI_DGNS_R primer region on the COI gene, the CytB_DGNS_F and CytB_DGNS_R primer region on the CytB gene, and the 18S_F4 and 28S_R8 primer region on the 18S and 28S gene. The primer sequence is compared to the nucleic acid composition of every major spider clade. The primer position in the gene is detailed below the primer name in brackets. The position in the gene is based on the alignment of the primer to the *Drosophila melanogaster* reference sequence. The COI and CytB gene sequences of all available spider species were obtained from the NCBI database. The 18S and 28S gene sequences of spider species were obtained from the NCBI database through the Wheeler et al. 2017 list. For better visibility, the sequences downloaded were merged into the major spider clades. The Araneomorphae - basal group contains species belonging to families Austrochilidae, Filistatidae, Gradungulidae and Hypochilidae. The bases S, H, W, Y, R and D in the primer sequence represent the IUPAC nucleotide codes for degenerate bases, which are common in universal primers. Note that the reverse primer is written in reverse complement, from 3’ to 5’. Figures were generated using the plot_alignments command in the R package PrimerMiner (Elbrecht & Leese 2017).

**Table 1.**
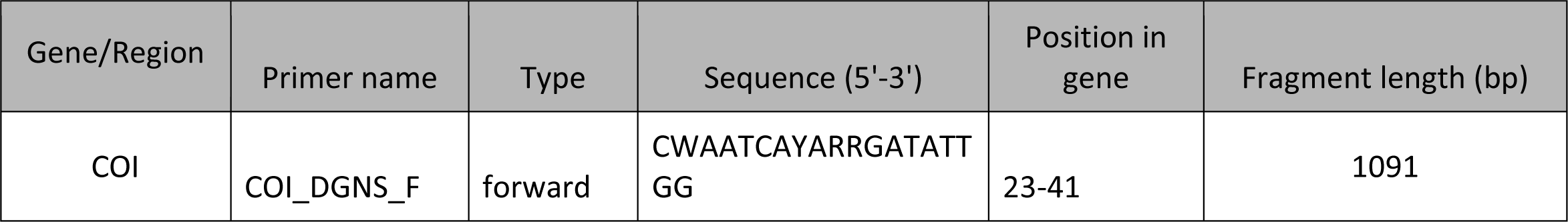
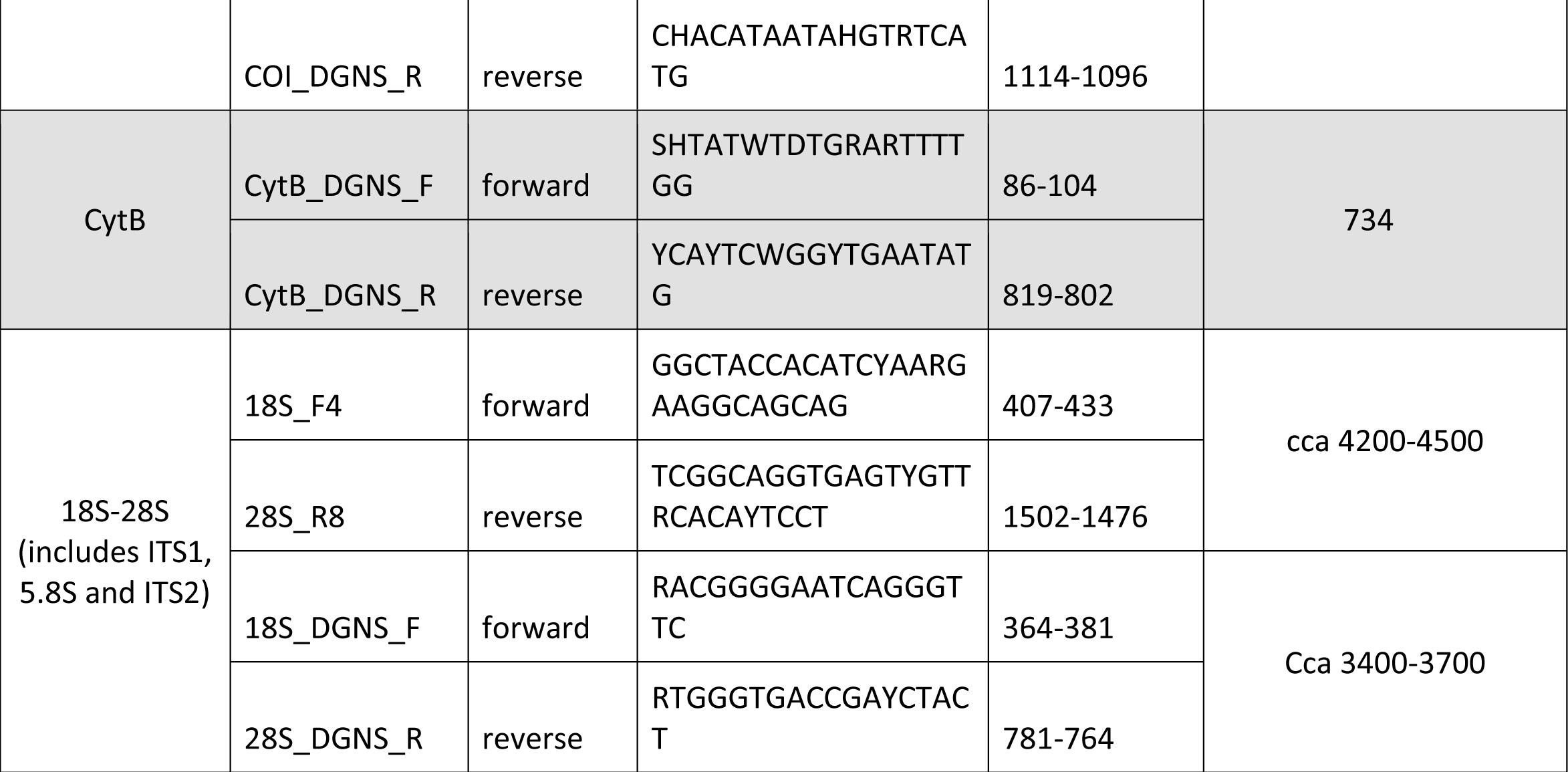
List of new barcoding primer sets established in this study. Shows the name of the primers, their orientation, sequence, position in the gene, and length of the amplified fragment, including primer-binding sites. The position in the gene is based on the alignment of the primer to the *Drosophila melanogaster* reference sequence.

Indexed PCR products were pooled in approximately equimolar amounts based on gel band strength. The mitochondrial and nuclear amplicons were pooled separately because, based on previous work, the shorter mitochondrial amplicons were expected to outperform the longer nuclear ones in library prep. Leftover primers were removed from the pools using 0.7X magnetic beads in accordance with the AMPure XP protocol (Beckman Coulter, Pasadena, CA). Nanopore library prep was then performed separately on each pool, using the SQK-LSK109 kit (Oxford Nanopore Technologies, Oxford, UK) following Pomerantz et al. (2022), and using the Short Fragment Buffer (SFB) for the mitochondrial and the Long Fragment Buffer (LFB) for the nuclear pool. Based on concentrations measured with the high-sensitivity dsDNA assay of the Qubit Fluorometer (ThermoFisher), libraries were merged into a final pool containing roughly 20 fmol of the mitochondrial and 30 fmol of the nuclear library. The higher amount of nuclear library was used because the shorter mitochondrial fragments were anticipated to sequence preferentially. The resultant pool was sequenced on a MinION using an R9.4.1 flow cell (Oxford Nanopore Technologies) following the manufacturer’s protocol. Sequencing was allowed to run for 72 hours to maximize read depth per sample.

### 2.4 Sequence processing

Sequences were processed following Pomerantz et al. (2022), with a few adjustments to the script. Samples were first demultiplexed by index combination using *minibar* (Krehenwinkel et al., 2019), and then split by barcode (COI, CytB or ribosomal cluster, based on inner primer sequences) using *NGSpeciesID* (Sahlin et al., 2021).

*NGSpeciesID* was also used to generate consensus sequences for each sample. Due to the co-amplification of contaminants (e.g. fungi), paralogs or NUMTs (Lopez et al., 1994), a single sample often yielded multiple consensus sequences. Sequences were therefore checked by BLAST search (Altschul et al. 1990) against the NCBI nucleotide database (accessed 10/2022), and only the correct sequence for each sample was retained. The whole script for processing of sequences using our primers is available in the supplementary material.

Sequences of COI and CytB were aligned in MEGA11 (Tamura et al., 2021) based on their reading frame, which was checked by translating the nucleotides into amino acids using the Invertebrate Mitochondrial code. For the 18S-28S amplicons, sequences were aligned with ClustalW MSA with a gap opening penalty of 15.00 and gap extension penalty of 6.66 in MEGA11 (Tamura et al., 2021). Alignments were visually inspected for obvious alignment errors, which were corrected by hand. The final concatenated alignment of all barcodes was 6121 bp long [CytB: 696 bp, COI: 1054 bp, and 18S-28S: 4371 bp (18S: 1436 bp, ITS1: 625 bp, 5.8S: 185 bp, ITS2: 470 bp, 28S: 1655 bp)].

As hyper-variable and misaligned sites can misinform phylogenetic inference, such sites (concerning some fractions of the ITS regions) were removed using Gblocks sequence editing (Castresana, 2000) with medium settings (b1 = 0.5, b2 = 0.5, b3 = 6, b4 = 6, b5 = ’All’). The resulting matrix comprised 5047 positions (82 % of the original matrix).

### 2.5 Phylogenetic inference

For phylogenetic inference, the concatenated alignment was partitioned into the three codon positions of the COI and CytB partial genes, and the ribosomal cluster was partitioned into 18S, ITS1, 5.8S, ITS2 and 28S, which were assumed to evolve at disparate rates and substitution models (see Tab. A.3 for best partitioning scheme as determined using ModelFinder (Kalyaanamoorthy et al., 2017) in IQTree). Phylogenetic inference was performed with IQtree (Nguyen et al., 2015) using the W-IQ-tree web application (Trifinopoulos et al., 2016) with ultra-fast bootstrap (ufb) (Minh et al., 2013) and SH-aLRT branch tests (Guindon et al., 2010) with 1,000 replicates each to assess branch support. Phylogenies were plotted with FigTree (Bouckaert et al., 2014).

### 2.6 Intraspecific and intrageneric genetic distance comparison

As our primers produce amplicons of longer length than commonly used barcoding markers (e.g. COI-5P), we wanted to assess whether there would be any difference in intrageneric genetic distances of our amplicons compared to the standard COI barcode traditionally obtained by Sanger sequencing. Therefore, using the NCBI nucleotide database (Sayers et al., 2022), we downloaded the first 658-bp COI barcode sequence found that corresponded to each species used in our case study. We only used genera represented by more than one species in our data set, allowing us to measure sequence distances between species within each genus (Tab. A.5 and A.6). We focused only on the COI marker due to the poor representation of CytB and the 18S-28S region for spiders in the NCBI database. After downloading the short fragment sequences, we aligned them in MEGA11 (Tamura et al. 2021) based on their reading frame as described above. After correction, a distance analysis was performed using the Kimura 2-parameter model (K2P) (Kimura, 1980) in MEGA11 (Kumar et al., 2016). Statistical analysis was performed in the R environment—version 4.2.0 (R Core Team 2021). To assess if there was any difference in the overall intrageneric genetic distance distribution of the investigated species between longer and shorter marker, we used general linear models and performed an ANOVA test on our two levels of the explanatory variable (short vs long COI marker; Pekár & Brabec, 2016).

Additionally, we wanted to evaluate the taxonomic resolution of our three barcoding markers combined (COI + CytB + 18S-28S) to just our COI barcode alone. For that, we used the final alignments of the COI sequences and the final alignment of all three barcodes combined (the alignment used for phylogenetic inference). We again performed a distance analysis using the Kimura 2-parameter model (K2P) (Kimura, 1980) in MEGA11 (Kumar et al., 2016), and correlated intraspecific and intrageneric distances of our three-marker set with the COI marker.

## 3. Results

### 3.1 Efficiency of PCR amplification

The Cytochrome B gene fragment was successfully amplified in 90.6 % (N = 129) of specimens, the COI fragment in 95.3 %, and the ribosomal cluster in 75.8 % (Tab. A.4). The unsuccessful amplifications represented a random mix of taxonomic identities. In many instances where we had several specimens from the same species, some of them successfully amplified while others did not.

### 3.2 Phylogenetic inference

Phylogenetic inference, obtained by combining all three markers and using maximum likelihood, produced a tree with high bootstrap values for most nodes, with all species and genera recovered as monophyletic (Fig. 2, Fig. A.1). Nearly all families and known higher clades, such as the oval-calamistrum clade, Araneoidea, marronoid clade, and the RTA clade (Wheeler et al., 2017; Fernández et al., 2018; Azevedo et al., 2022; Kulkarni et al., 2022;), were also recovered as monophyletic. Phylogenetic inference was enhanced by cleaning the alignment with Gblocks, which removed the most variable, saturated and/or poorly aligned sections of the ITS regions. The resulting phylogeny after Gblocks cleaning recovered Dionycha as a monophyletic clade (Fig. 2) and exhibited increased node support on average. On the other hand, the phylogenetic inference based on only the COI barcoding region produced a tree that does not support any of the known higher clades, in some cases even families (e.g. Agelenidae; Fig. A.2).

**Figure 2.**
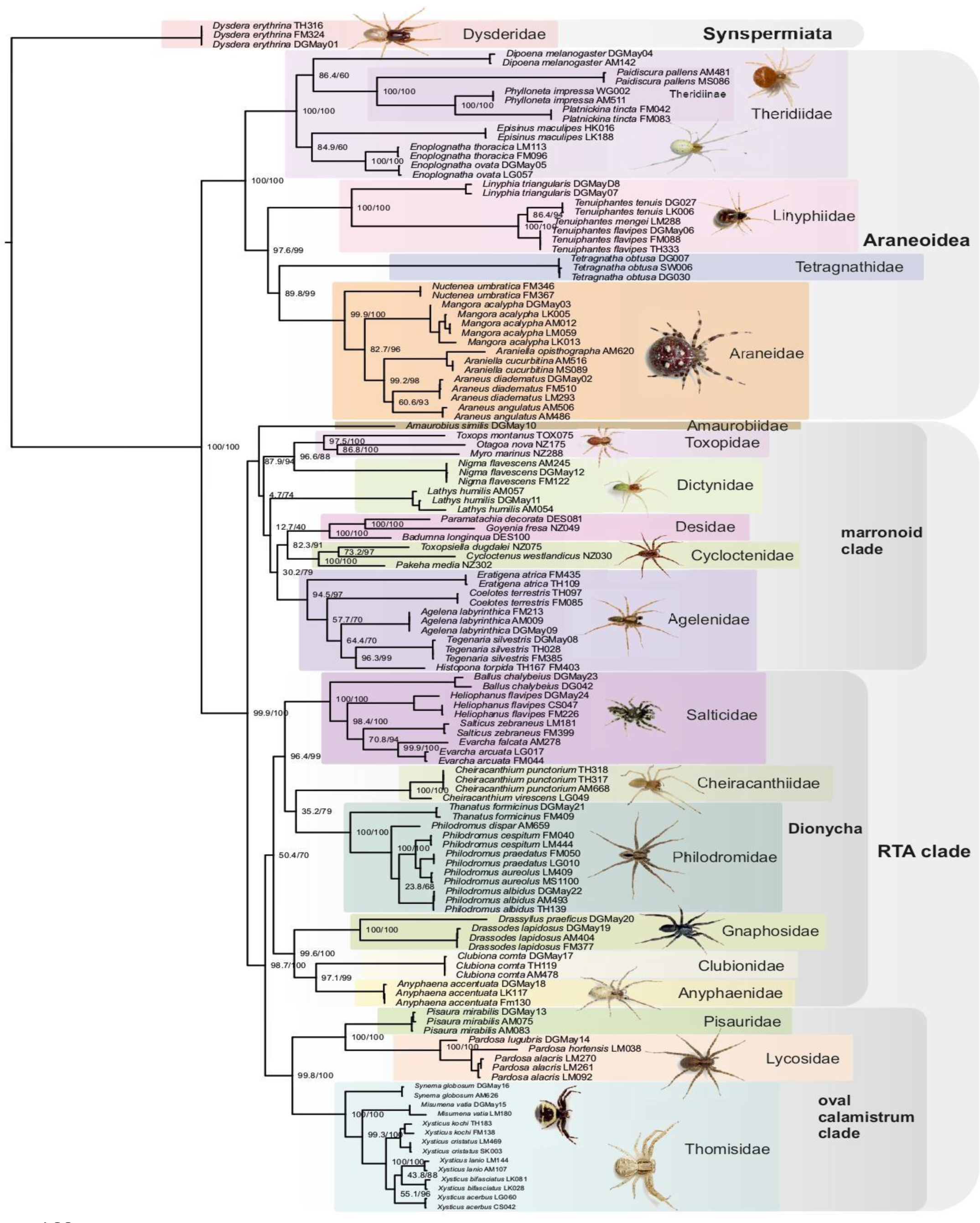
Maximum likelihood phylogeny inferred from the Gblocks-cleaned and partitioned alignment of the three amplicons obtained using the newly designed primers COI_DGNS_F/R and CytB_DGNS_F/R, and the published primers 18S_F4 / 28S_R8 (Krehenwinkel et al. 2019). Major clades widely recognized in spider systematics and recovered in previous multi-locus approaches are marked with color and labelled. Numbers at nodes give the ufb- and SH-aLRT node support values. The two specimens of *Histopona torpida* (Agelenidae) were merged into one sequence because one specimen only recovered the mitochondrial and the other only the nuclear amplicon.

### 3.3 Intraspecific and intrageneric genetic distance comparison

The difference in the distribution of intrageneric genetic distances between the long and short COI marker was not significant (LM, F1,54 = 0.262, P = 0.611; Fig. 3). The change in kimura2 distance value from short marker to long marker for each respective species comparison was random, and ranged from -2.57 % to 1.86 %, but the median value was always around 10 % for both markers. The data showed a few genetic distance comparisons that were recognized as outliers from an otherwise normal distribution of distance values. For the lower distances, these were species that are recognized as recently diverged (e.g. *Philodromus* species from the Aureolus group: *P. cespitum*, *P. aureolus* and *P. praedatus*); for higher distances, these were species belonging to Theridiidae and Araneidae (Figure 3 ; Tab. A.5 and A.6).

**Figure 3.**
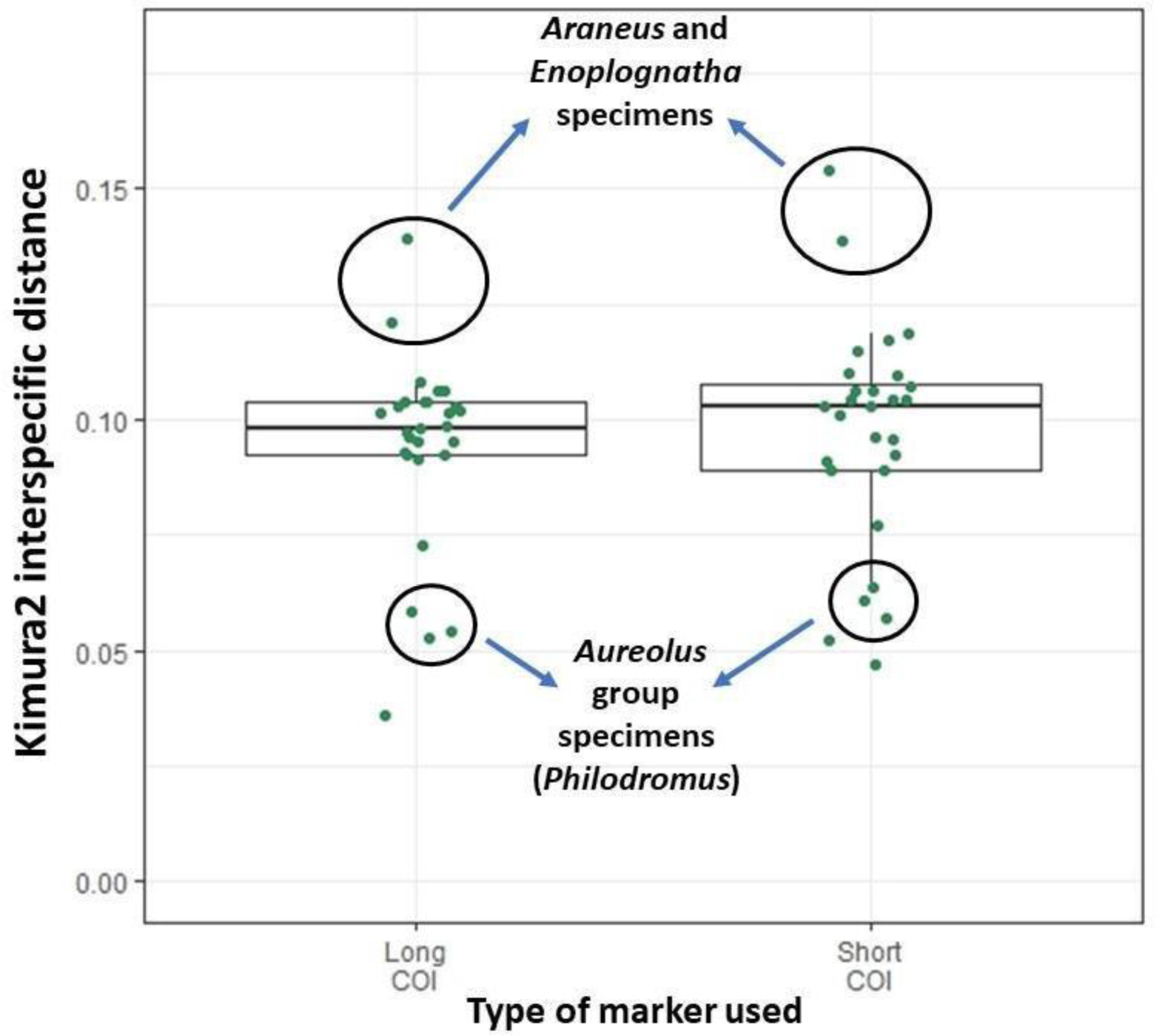
Comparison of intrageneric kimura2 distance distribution between the long COI fragment obtained by our newly designed primers and the common COI-5P barcode. The dots inside the boxplots represent individual intrageneric genetic distance values.

We found a significant correlation of pairwise genetic distances between our long COI marker and all the three long markers combined (Linear regression, p < 0.001; Fig. A.3). However, the three markers combined didn’t show a perfect association with COI. An overlap between intraspecific and intrageneric distances was present in the combination of all three barcodes together. But there was a visible genetic distance gap between intraspecific and intrageneric distances in the COI barcode alone, where intraspecific samples went up to 2.5 %, and intrageneric samples only started at 5.6 % genetic distance.

## 4. Discussion

### 4.1 Efficiency of PCR amplification

Our primers for COI and CytB worked in more than 90 % of cases, while amplification of the ribosomal cluster was successful in 75 % of specimens. For the nuclear markers 18S_F4 and 28S_R8 (taken from Krehenwinkel et al., 2019), the unsuccessful outcome occurred in a random assortment of families. In many instances where we had several specimens from the same species, some of them successfully amplified while others did not. Therefore, primer amplification bias should be minimal, as additionally supported by our in-silico evaluation of primers (Fig. 1). A possible cause of the lower success rate for the nuclear amplicon could be lower DNA integrity for some samples, as the DNA strands might have already been broken into fragments smaller than ∼4500 bp. The specimens we collected into absolute ethanol were stored for several months at room temperature, which might have caused some extent of DNA degradation (Krehenwinkel & Pekar, 2015; Marquina et al. 2021). However, for all of these specimens, we managed to amplify at least one of the mitochondrial markers, indicating that DNA fragments of approximately 1 kb were still intact.

This is why, in case of difficulty amplifying the ∼4,500 bp fragment, we propose another primer pair that we designed for the amplification of the 18S-28S region. The primer pair 18S_DGNS_F / 28S_DGNS_R produces a ∼ 3,500 bp fragment (Tab. 1; Fig. A.4). These newly designed primers were found to be successful in in-silico tests for the whole spider clade (order Araneae) and in-vitro tests on the 24 spider species used in the PCR optimizations. Because the resultant amplicon overlaps with roughly 3,000 bp of the one obtained from primers 18S_F4 and 28S_R8 (Krehenwinkel et al., 2019), one can easily combine data from specimens that worked with the larger fragment and with the smaller fragment, if needed. Furthermore, the 28S_DGNS_R primer was designed to match spider templates while having mismatches with plants, fungi and nematodes (Fig. A.4), thus suppressing the amplification of these potential contaminant taxa. Such unwanted taxa have been shown to preferentially amplify when the length of their nuclear ribosomal cluster is shorter than that of the target taxa (Wattier et al. 2002; Krehenwinkel et al., 2019).

### 4.2 Phylogenetic inference

Our test showed that the combination of our three long amplicons exhibited high phylogenetic informativeness on both shallow and deep evolutionary time scales, while this was not the case when using the COI amplicon alone. Our approach successfully recovered the major monophyletic clades recognized in spider systematics, such as Araneoidea, the RTA clade, the marronoid clade, the oval-calamistrum clade and Dionycha (Azevedo et al., 2022; Fernández et al., 2018; Kulkarni et al., 2022; Wheeler et al., 2017; but note that the latter was only recovered after removing misinforming hypervariable sections from the ITS-regions using Gblocks). Dictynidae was paraphyletic, which has recently been indicated independently by other studies using different approaches (Crews et al., 2020; Kulkarni et al., 2022). All other families, genera and species were recovered as monophyletic. The only significant discrepancy to the current state of the art in the higher-level phylogenetics was the position of Tetragnathidae (Kulkarni et al., 2022), which might rather be an effect of the taxonomic sampling than the method. We note that the taxonomic sampling used for this study was mainly guided by the aim to test the protocol on representatives of a broad variety of spider families, and not to sample evenly across the whole spider tree of life. Hence, we lack information on some groups (e.g. Mygalomorphae), especially for the deeper and intermediate nodes.

Generally, intermediate nodes (i.e., relationships between families) had mixed node support, with particularly low bootstrap values in some nodes that are known to be hard to resolve (Azevedo et al., 2022; Kulkarni et al., 2020). This is probably due to heterogeneous evolutionary rates throughout time. Nevertheless, the results show that the approach is highly suitable, especially for rapid and cost-efficient phylogenetic inference on the (sub-)family or genus level, or to inform about taxonomy, systematics or comparative analyses. If the aim is to generate a broad-scale phylogeny to enhance the accuracy of higher-level topologies, we recommend using a phylogenetic backbone constraint informed by published genomic/transcriptomic analyses during phylogenetic inference (see, e.g. Wheeler et al. 2017; Wolff et al., 2022).

All things considered, the inferred phylogeny was overall consistent with the results from approaches using hundreds to thousands of loci (Azevedo et al., 2022; Fernández et al., 2018; Kulkarni et al., 2022) while requiring only a fraction of the labour and cost needed for such approaches. Specifically, our approach takes approximately one to two weeks of hands-on lab work and currently costs roughly 5 € per specimen, a fraction of the price for ddRAD sequencing (approx. 40 - 60 € per specimen if the whole Illumina sequencing lane is used), UCE sequencing with targeted enrichment (approx. 100- 120 € per specimen if the whole Illumina sequencing lane is used), or genome sequencing (prices vary depending on the coverage, but usually costs are over 1000 € per specimen).

Our approach is generally useful for researchers that lack access to high-state-of-the-art molecular laboratories that are set up for genome-level taxonomic analysis.

Everything can be performed with equipment available in common molecular laboratories or even in improvized laboratories (e.g. a portable laboratory in the rainforest; Pomerantz et al. 2018) if an ONT MinION is provided. Furthermore, our approach can be useful for researchers who want to process large collections of taxonomically complex species mixtures simultaneously and obtain not only barcodes for taxon assignment but the phylogenetic inference. Results for such large and complex species mixtures are almost unattainable with approaches using thousands of loci as they are costly and produce immense amounts of data that are computationally intensive. On the other hand, as mentioned before, processing such complex sample sizes with standard COI barcodes alone is insufficient. Lastly, our protocol could be useful even for smaller barcoding projects, but with the help of the Flongle Flow Cell (Oxford Nanopore Technologies). The Flongle Flow Cell produces up to 2Gb of output at the cost of about 90 €, making it an inexpensive alternative to the MinION’s standard flow cell when one wants to process a small sample size.

### 4.3 Intrapecific and intrageneric genetic distance comparison

COI barcodes are commonly used for estimation of genetic distances of arthropod species and consequently assist in species delimitation (Čandek & Kuntner 2015; Zhang & Bu, 2022). We found that the 1,054-bp COI marker obtained with our primers does distinguish between specimens of the same species and different species of the same genera when comparing genetic distances (Fig. A.3). Furthermore, we confirmed that our long COI marker does not significantly deviate in intrageneric genetic distance compared to the traditionally used 5’ COI region (COI-5P), and the median value is around 10 % for both (Fig. 3). We observed a trend of some genera having lower or higher intrageneric genetic distance than the median value, often deviating from the otherwise normal distribution. The divergence of specific spider families and genera from the median value has been shown before (Čandek & Kuntner, 2015), and our results also suggest, even with the small sample size, that a taxonomically universal barcoding gap threshold is impractical (Yassin et al., 2010). Thus, when using genetic distance for species delimitation in spiders, we recommend preparing intra- and interspecific genetic distances of many specimens of the same genera or family as a reference, and not relying on standard thresholds.

### 4.4 Conclusion

Here, we develop a simple, cost-effective and accurate approach for DNA barcoding of spiders, using long mitochondrial and nuclear ribosomal amplicons, totalling to >6 kb of data per specimen. By combining three unlinked loci with large amplicon sizes, we are able to scale up barcoding from simple taxon assignment to large-scale phylogenetic inferences of large and taxonomically complex species mixtures. This was accomplished via two PCRs (one nuclear and one mitochondrial multiplex) and a simple sequencing protocol that can be easily carried out with basic laboratory equipment, making it widely accessible to scientists without extensive resources. The phylogeny inferred using our protocol was overall consistent with results from approaches based on hundreds to thousands of loci, while requiring just a fraction of the cost and labor of such approaches. Though we demonstrate the efficacy of this approach in spiders, with just a few nucleotide adjustments, the primers presented here could be used on a broader range of arthropods and therefore be useful not only to arachnologists but to other arthropod taxonomists.

## Author contributions

**Domagoj Gajski:** Data curation; Formal analysis, Investigation, Methodology, Project administration, Software, Visualization, Writing - original draft; **Jonas O. Wolff:** Data curation, Investigation, Methodology, Visualization, Writing - original draft; **Anja Melcher:** Data curation, Investigation, Methodology, Writing - review & editing; **Sven Weber:** Data curation, Investigation, Methodology, Writing - review & editing; **Stefan Prost:** Conceptualization, Funding acquisition, Software, Writing - review & editing; **Henrik Krehenwinkel:** Conceptualization, Funding acquisition, Project administration, Supervision, Resources, Writing - review & editing; **Susan R. Kennedy:** Conceptualization, Data curation, Funding acquisition, Project administration, Investigation, Methodology, Resources, Software, Supervision, Writing - original draft.

## Supporting information

Script for sequencing data processing

## Acknowledgements

We thank Arno Grabolle, Guido Gabriel and Jim McLean for their permission to reproduce their photos in Fig. 2. Additionally, we thank Dr. Stano Pekár and Dr. David Ortiz for giving us advice on how to improve our manuscript.

## Funding sources

This study was supported by grants from the German Research Foundation (DFG) in the framework of the priority program SPP 1991: TAXON-OMICS KE 2647/1-1. DG is a Brno PhD Talent Scholarship Holder—Funded by the Brno City Municipality. Additionally, for participation in this project, DG was supported by the Operational Programme Research, Development and Education “Project Internal Grant Agency of Masaryk University” (No. CZ.02.2.69 / 0.0 / 0.0 / 19_073 / 0016943). JOW was supported by the Deutsche Forschungsgemeinschaft (DFG, German Research Foundation) Grant 451087507.

## Declaration of interest

none

## Abbreviations

UCE: ultraconserved elements
ddRAD: double digest restriction-site associated DNA
NCBI: National Center for Biotechnology Information
BOLD: barcode of life data
ITS: internal transcribed spacers
COI: Cytochrome Oxidase Subunit I
CytB: Cytochrome B

---

**Table A.1.**
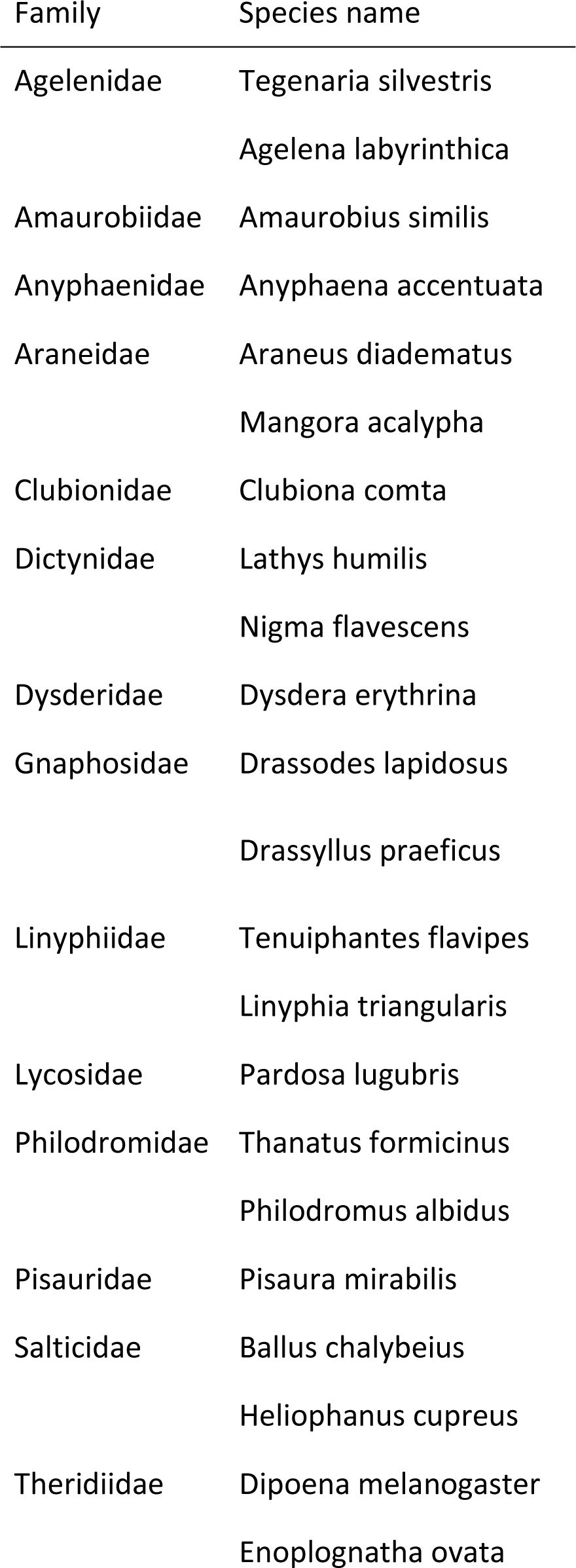
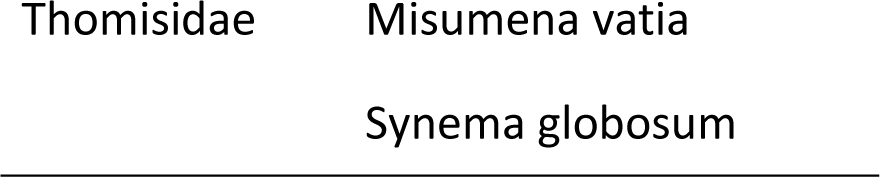
List of spider species used for the primary trial for the development and optimization of barcoding primers. Two specimens of *Linyphia triangularis* were sequenced; therefore, we have a total of 24 species.

**Table A.2.**
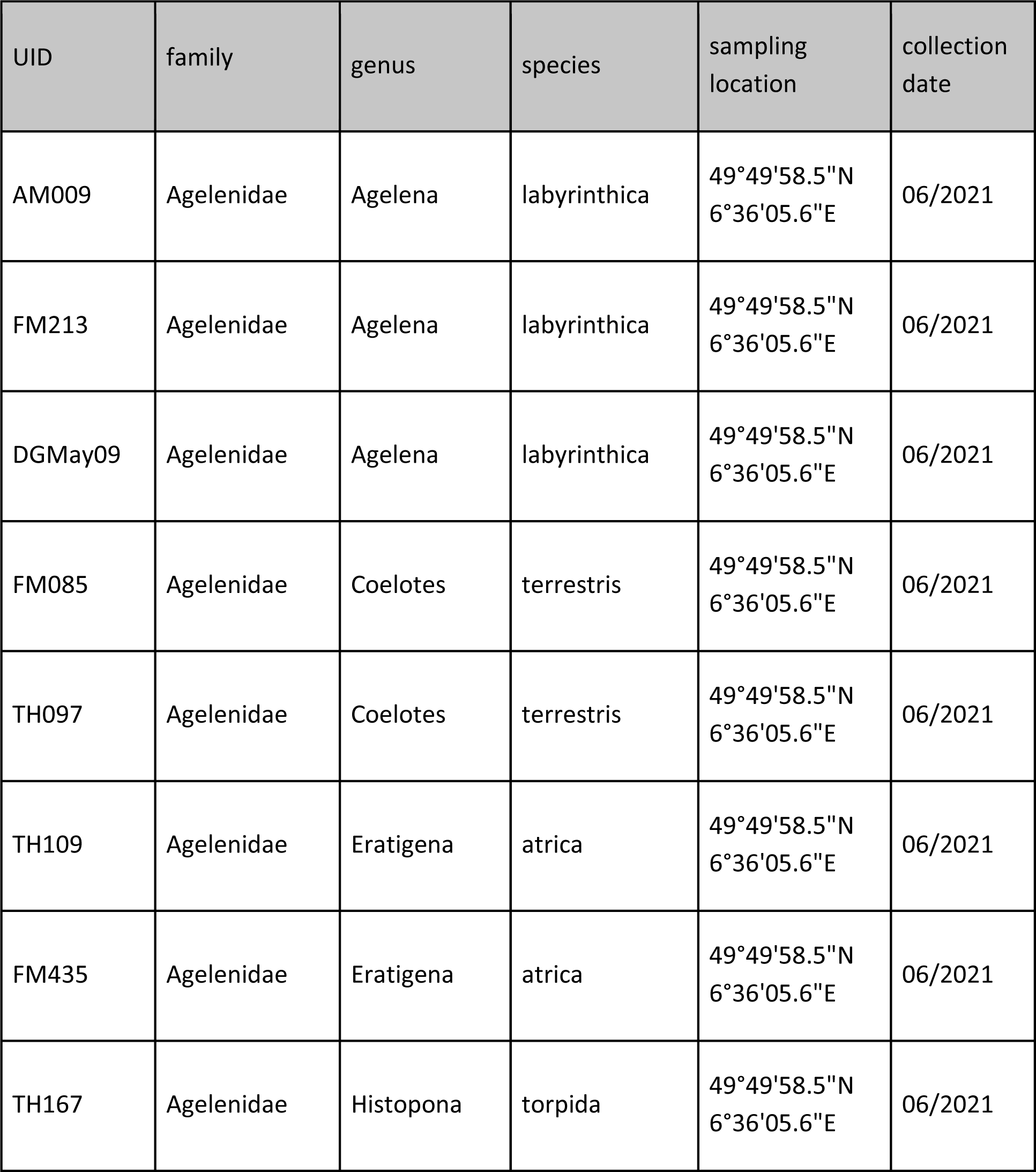
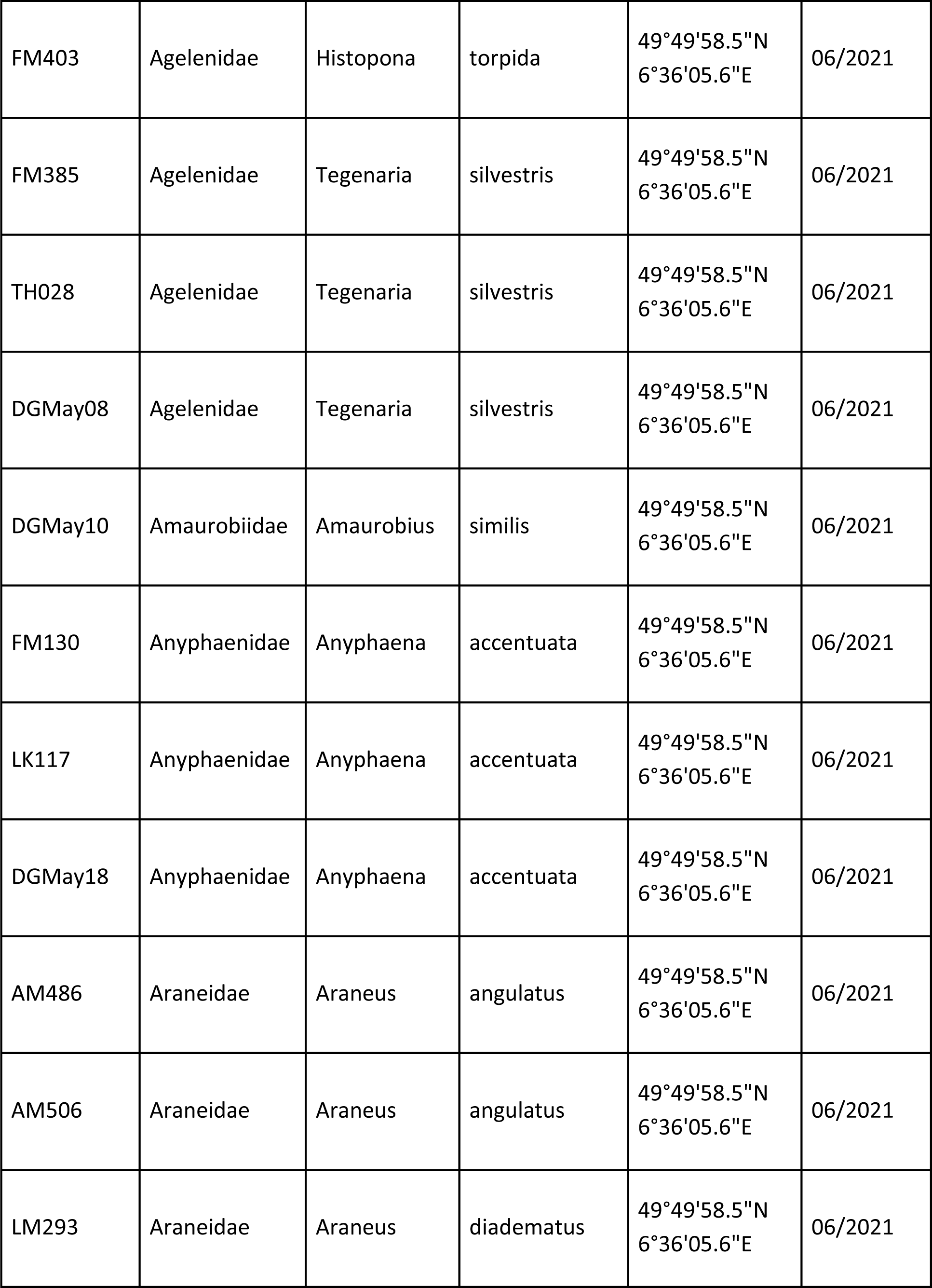
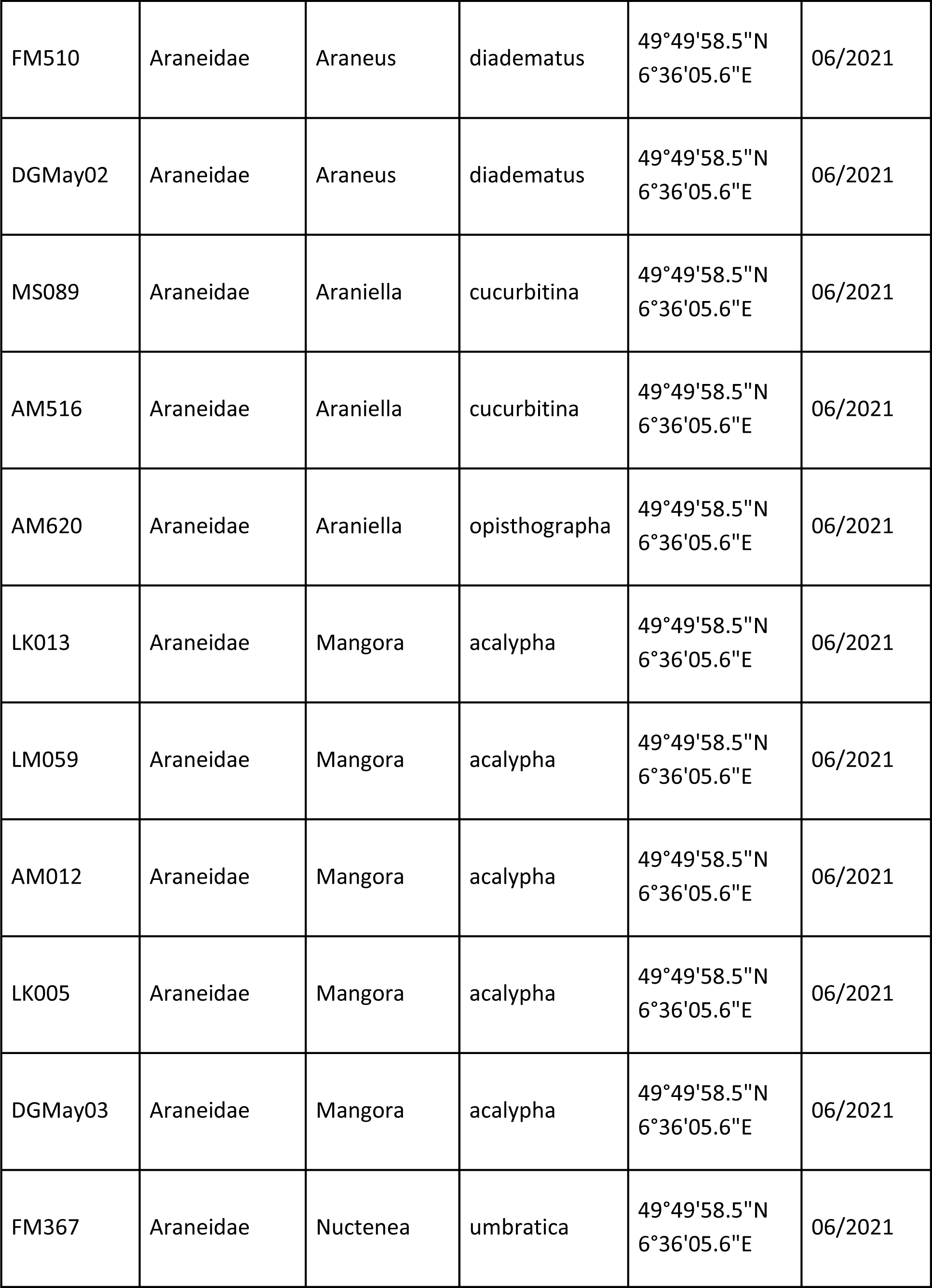
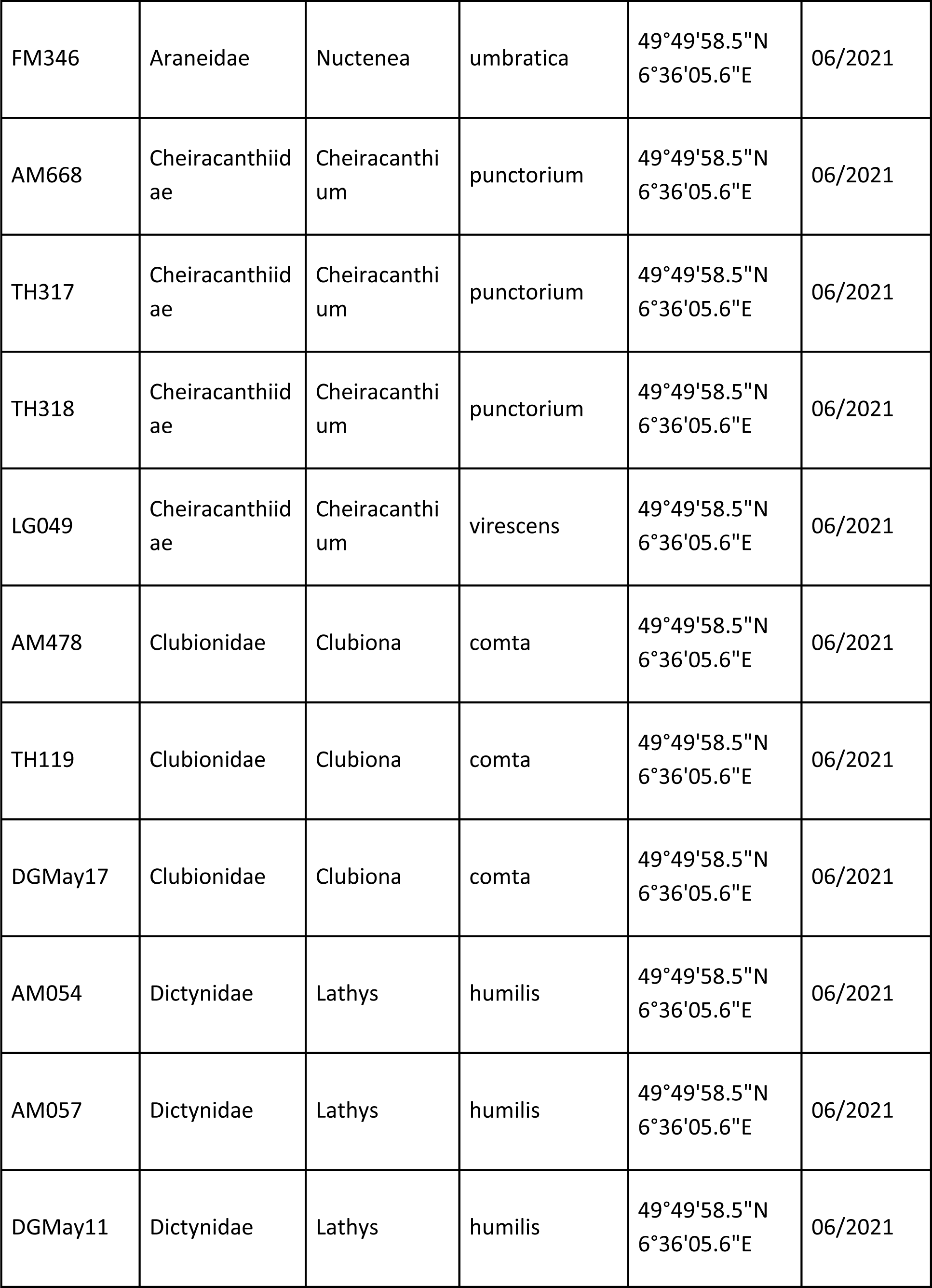
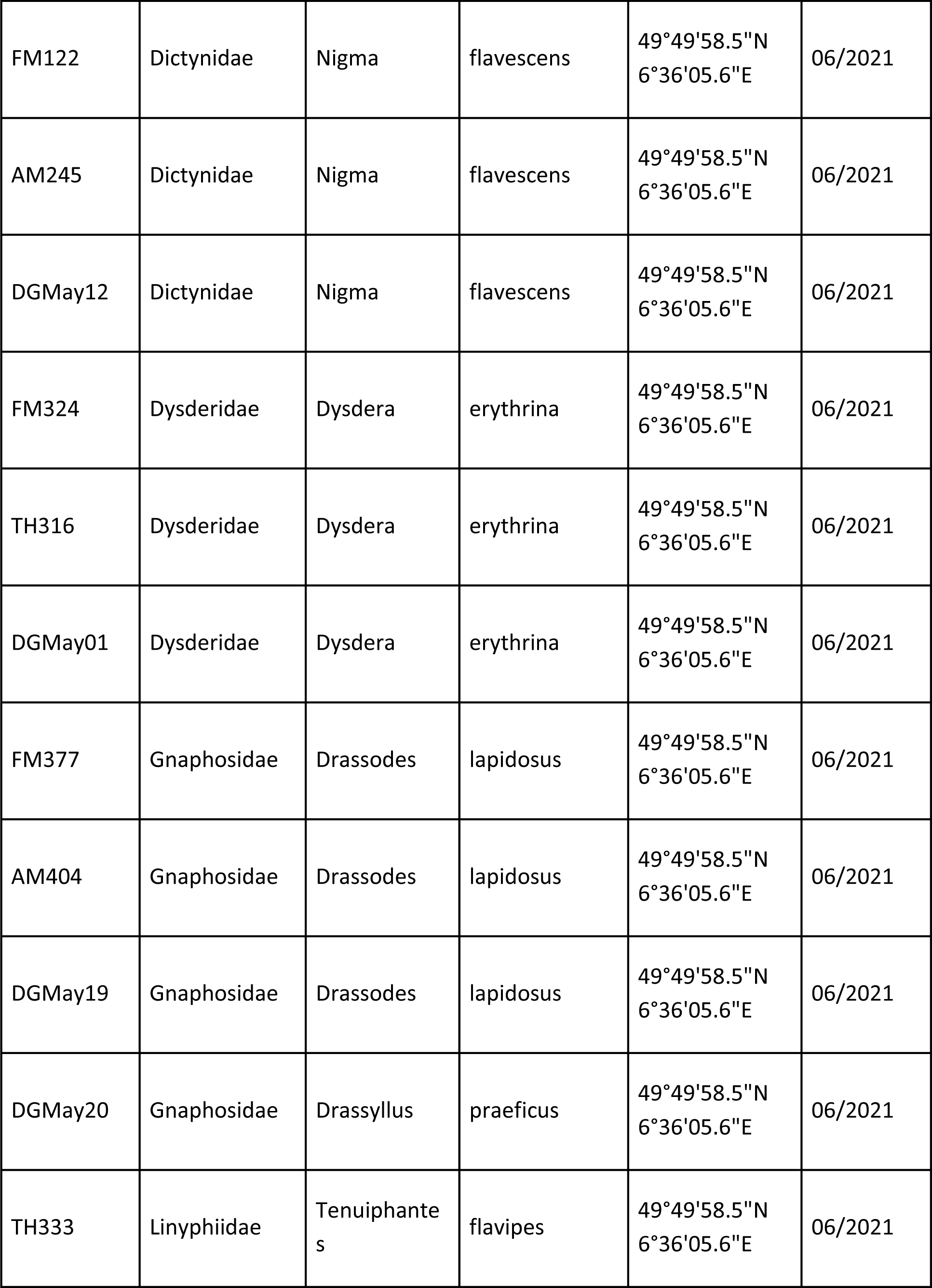
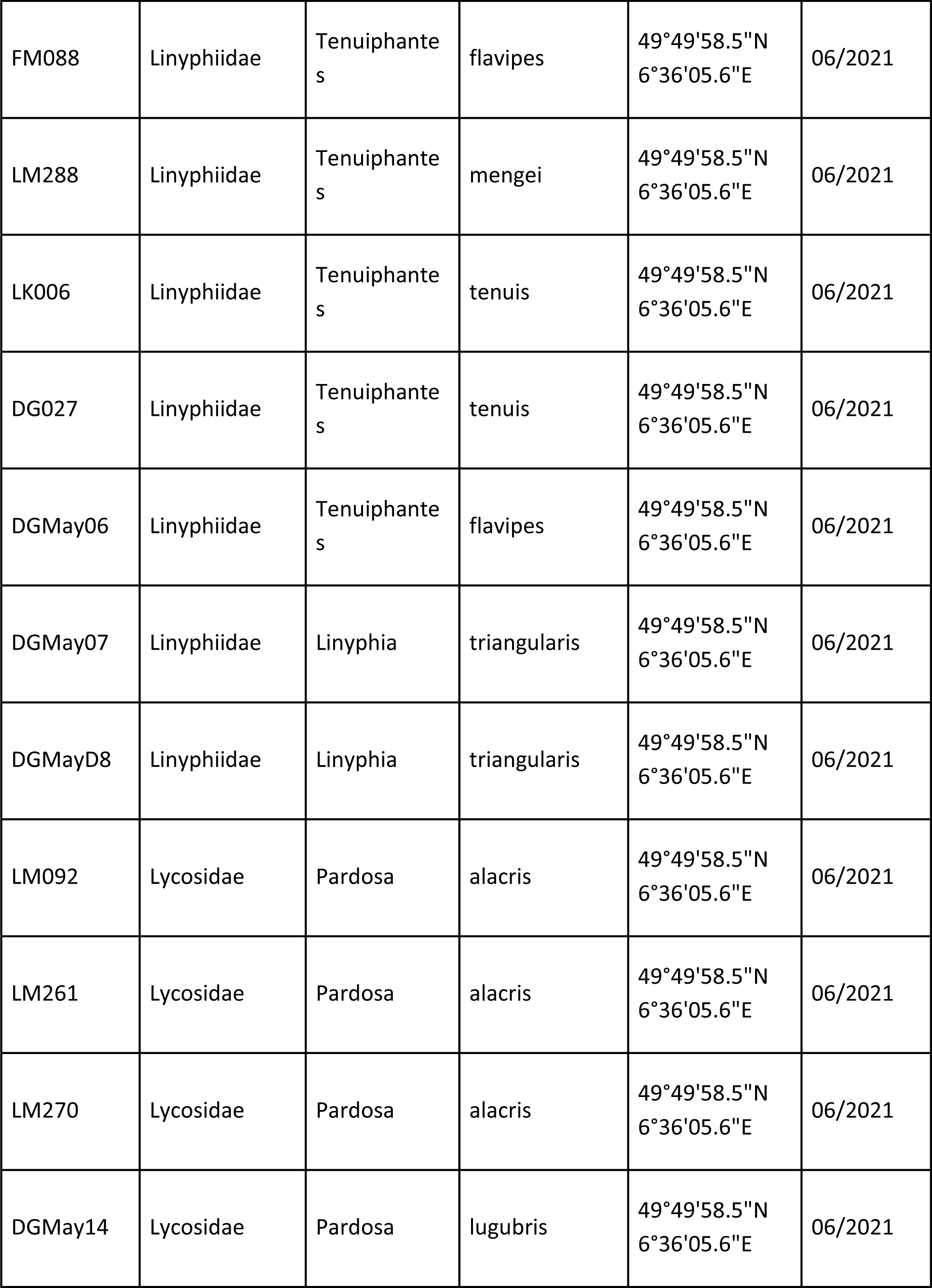
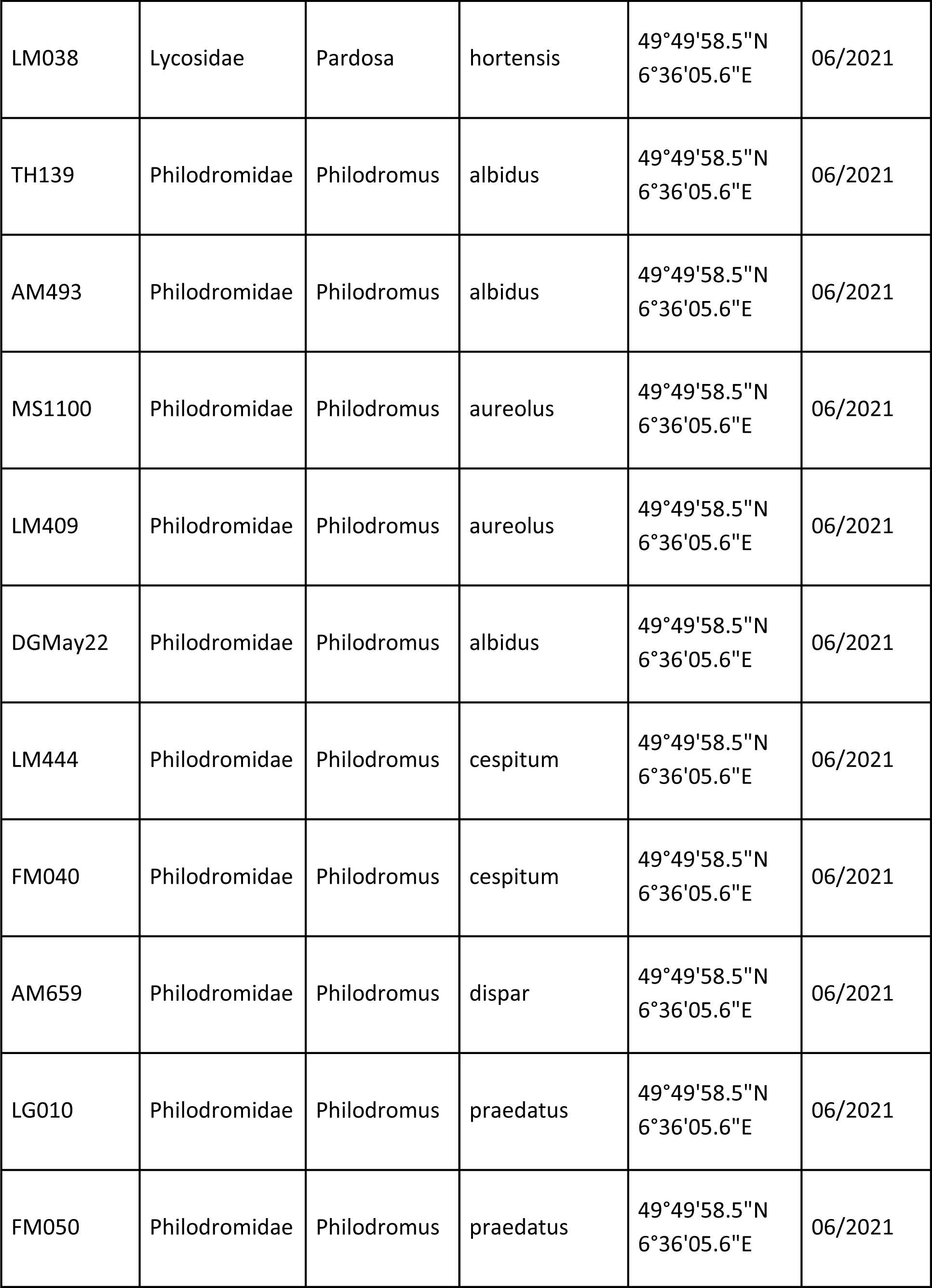
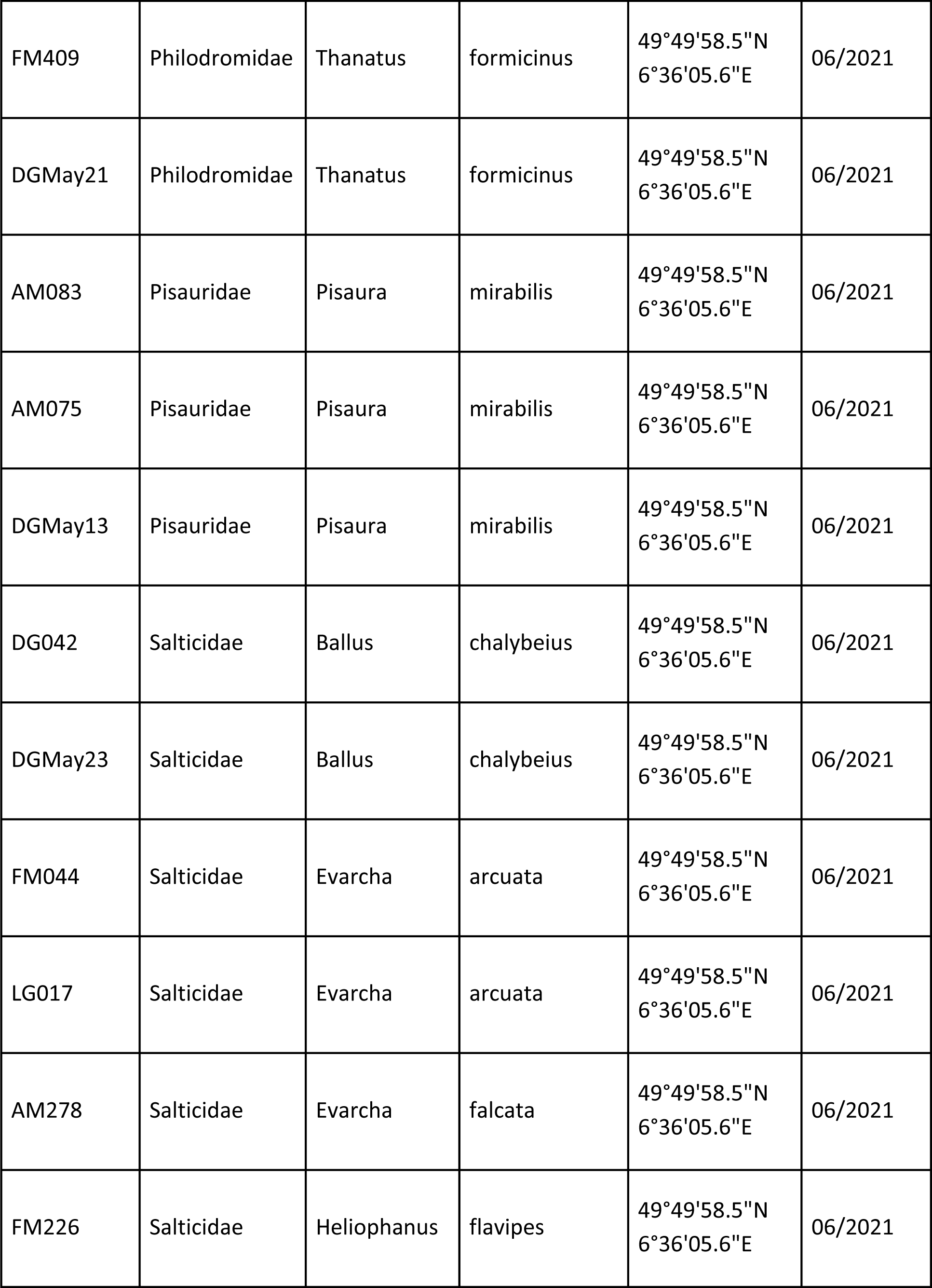
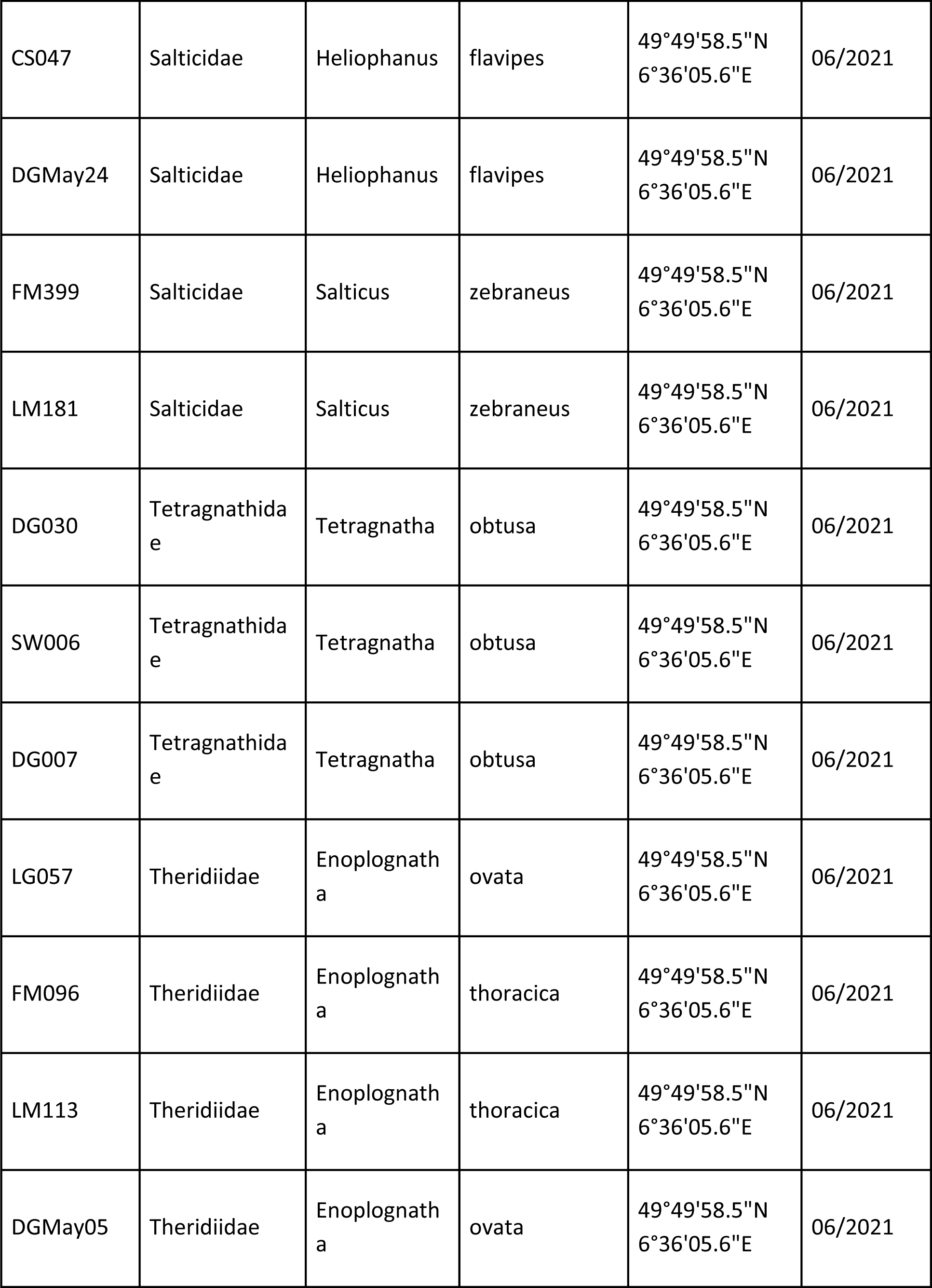
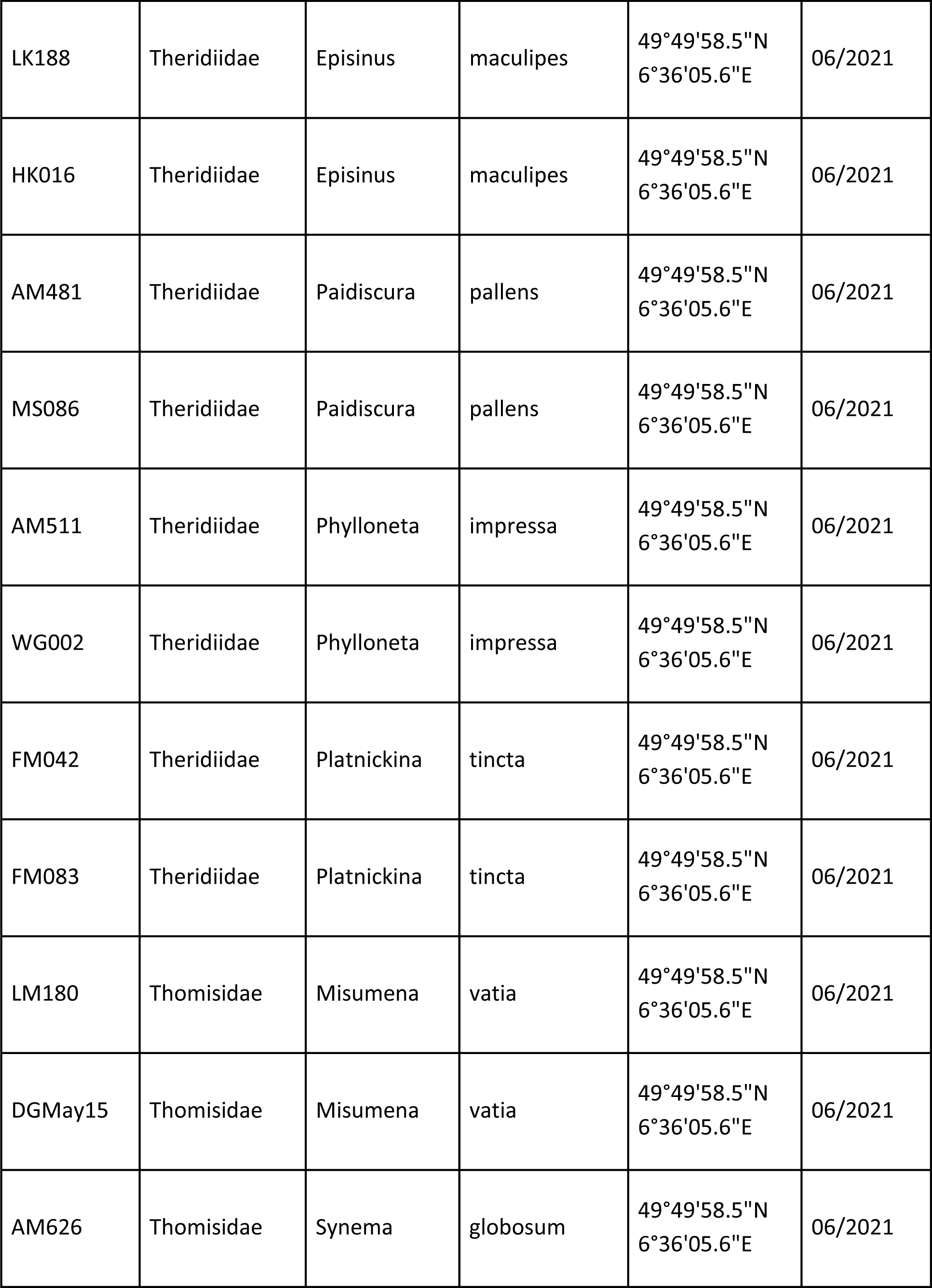
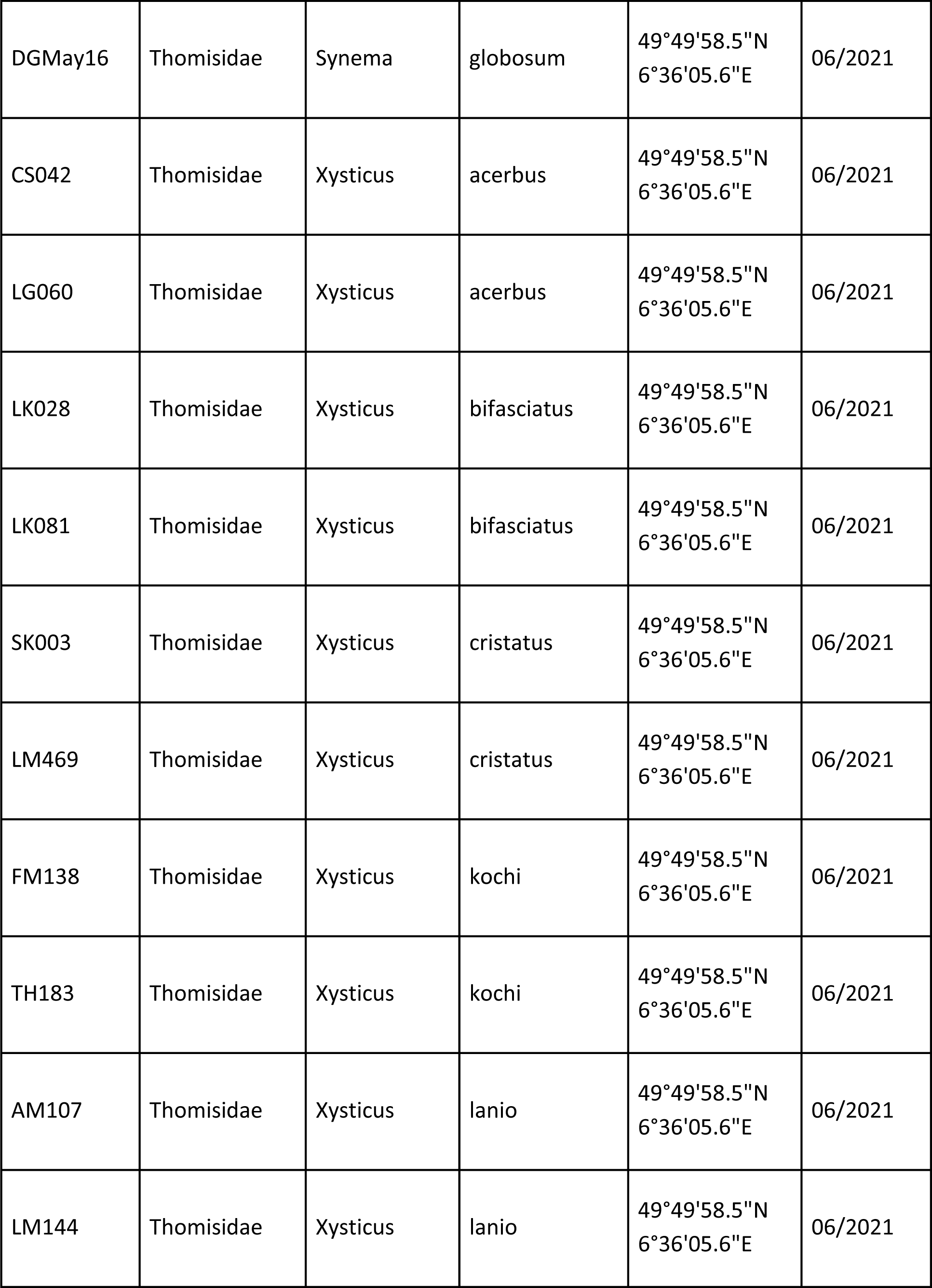
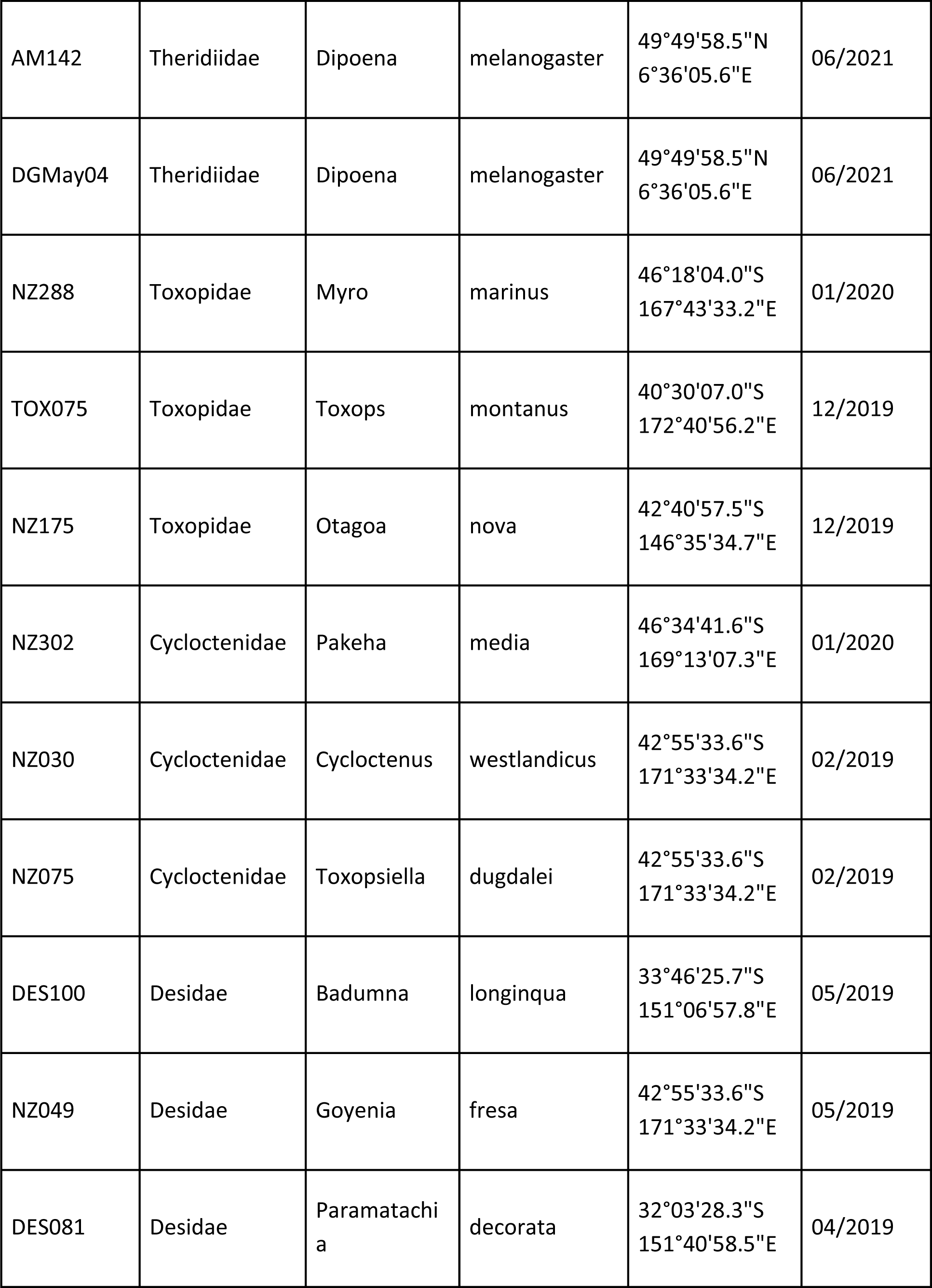
List of material with Genbank accession numbers (will be added upon acceptance of manuscript!). UIDs (“UID” = unique identifier) starting with “DGMay” were used in the PCR optimization trials; all others belong to the case study.

**Table A.3.**
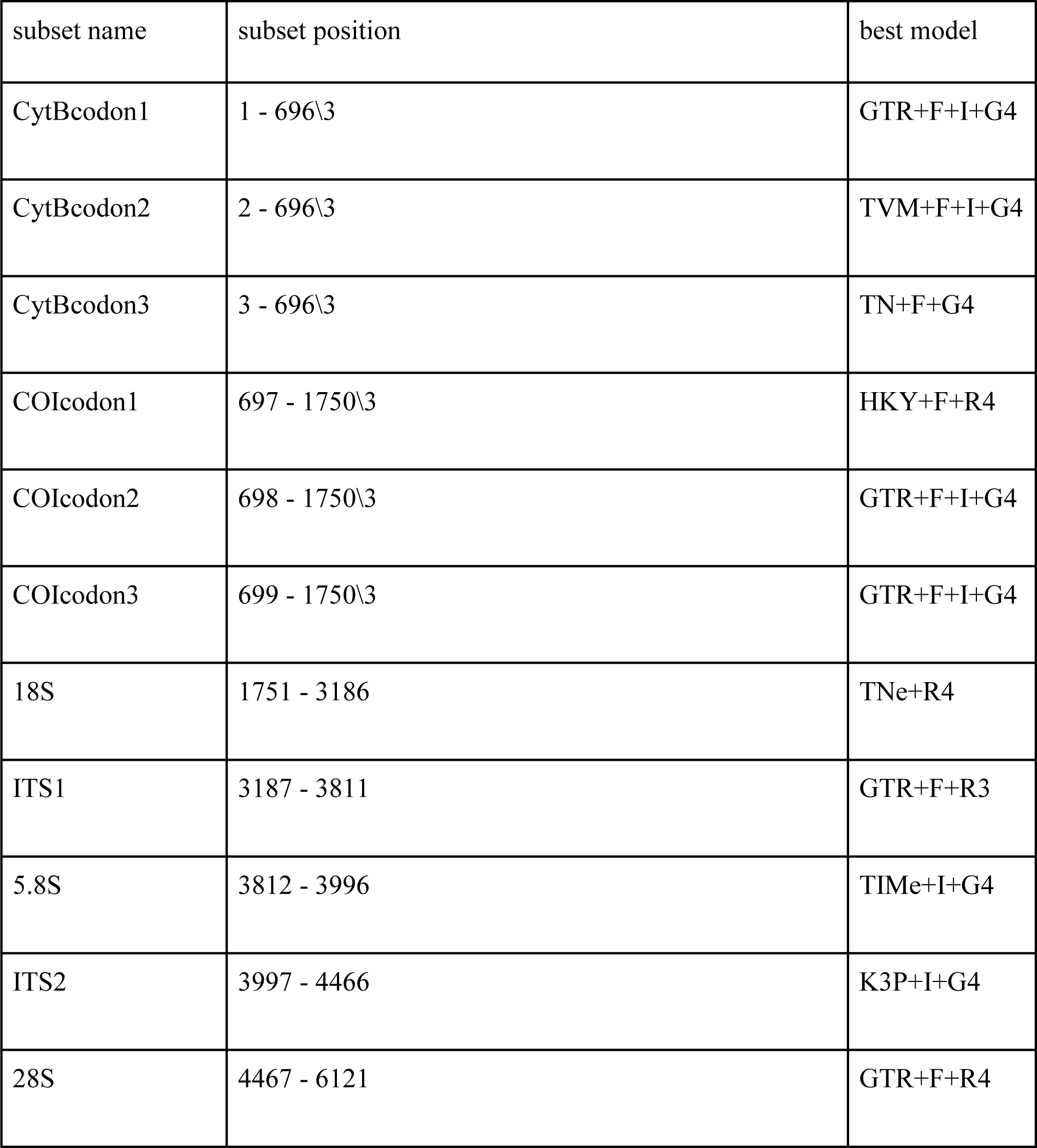
Best partitioning scheme and models used for phylogenetic analysis, as determined by ModelFinder (Kalyaanamoorthy et al., 2017) in IQTree (Nguyen et al., 2015).

**Table A.4.**
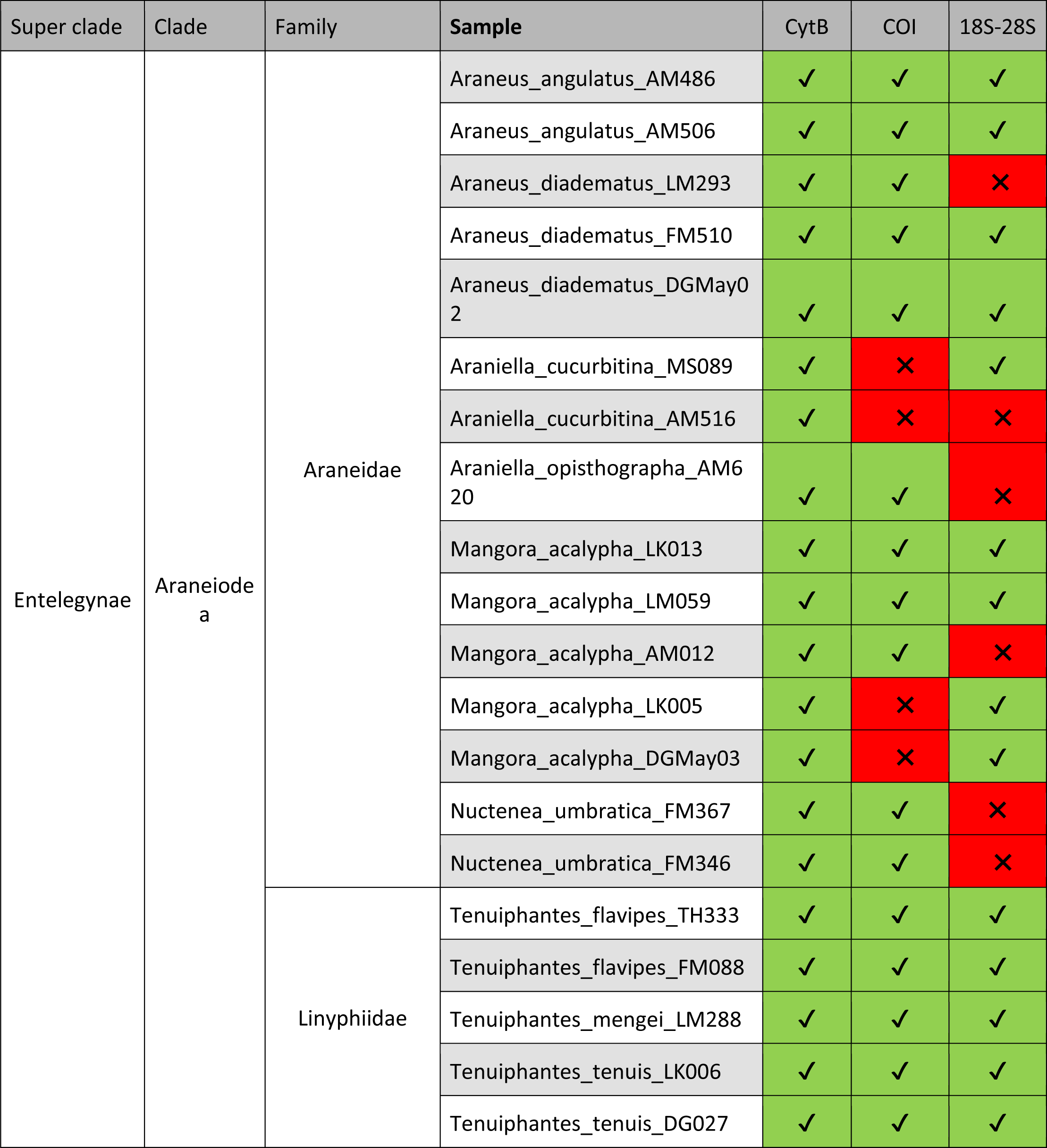
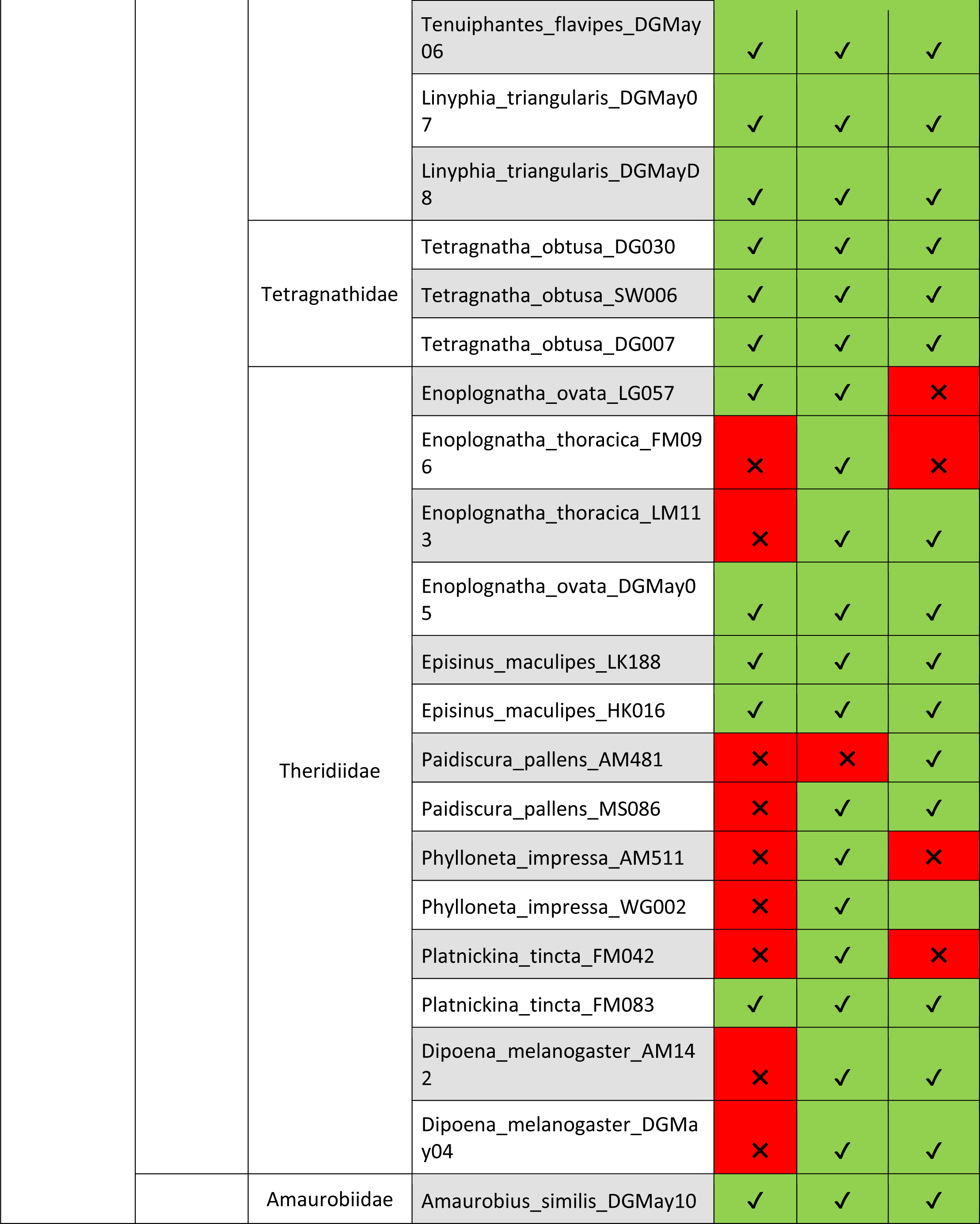
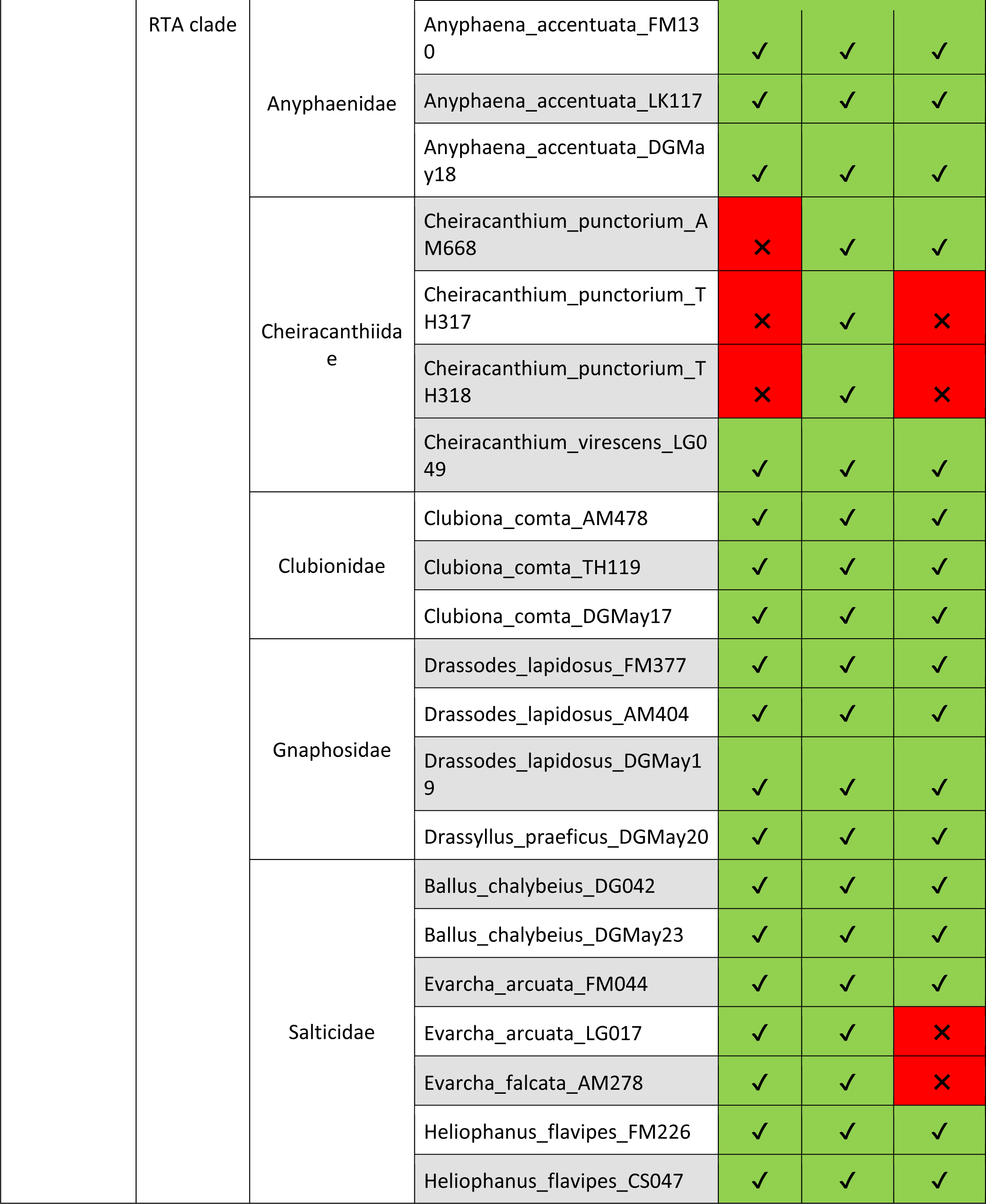
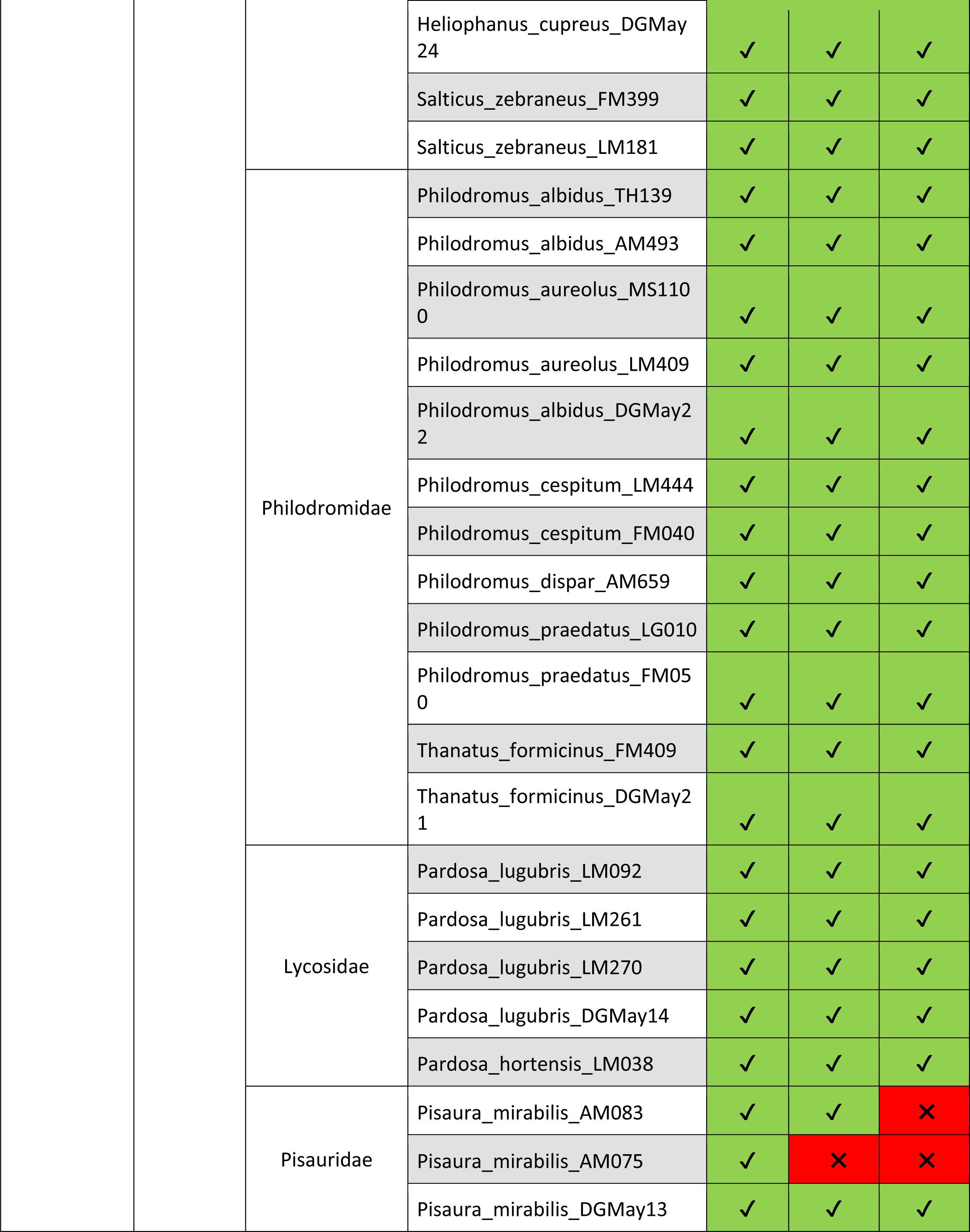
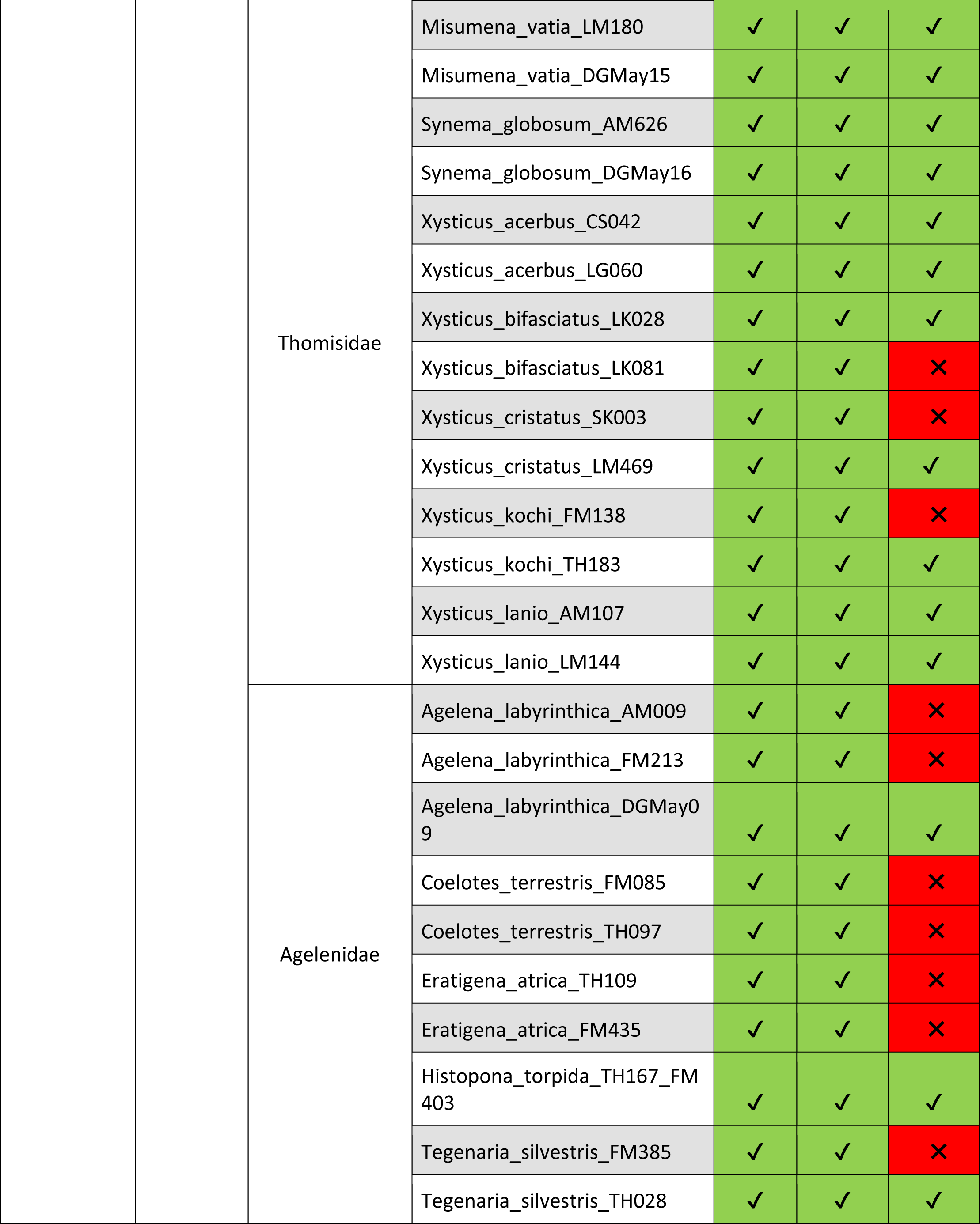
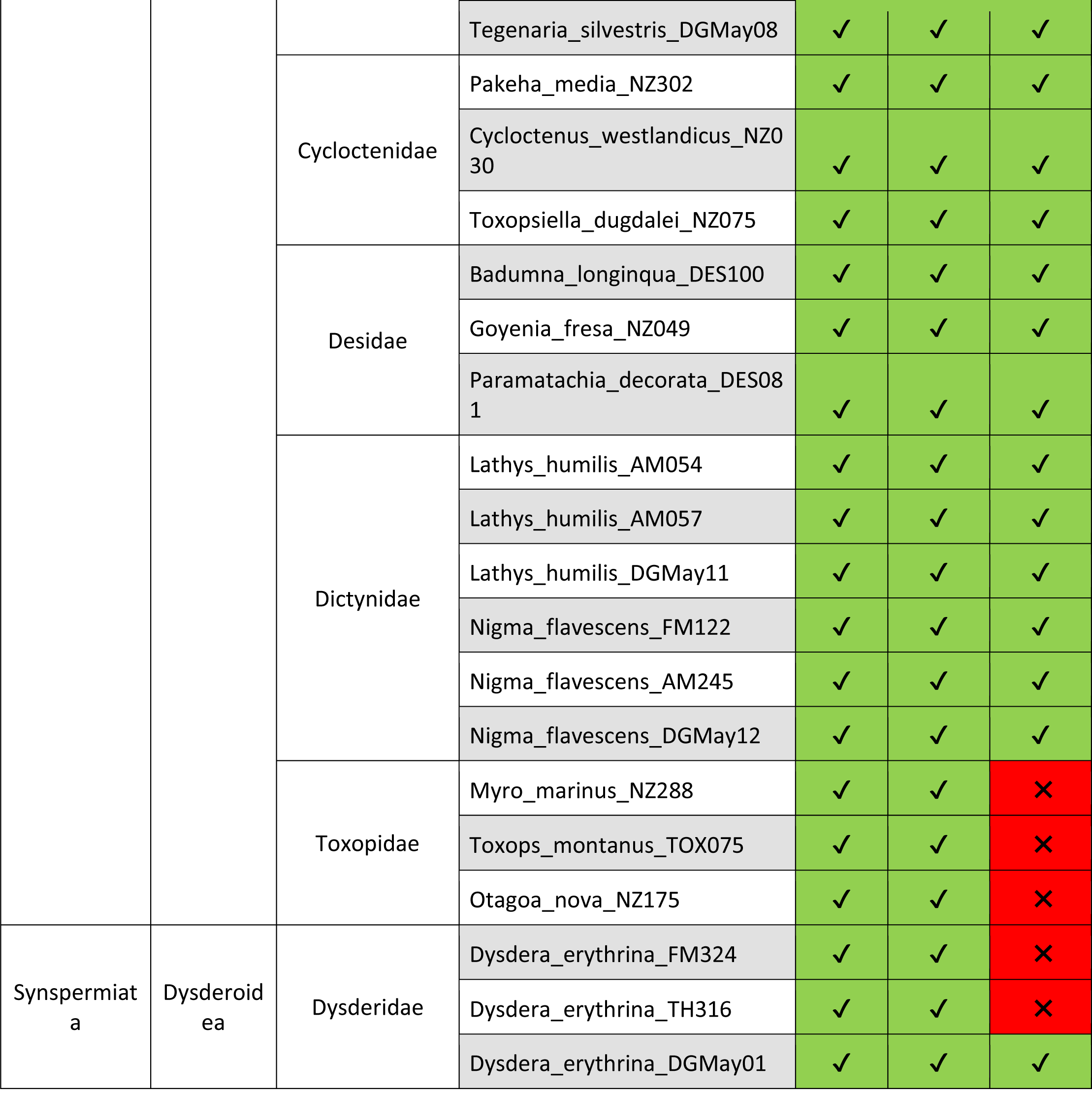
List of all samples used in our case study and success or failure of amplification and sequencing of target amplicons. Green/check mark denotes that the protocol successfully worked for a given amplicon for a given specimen, while red/X denotes failure to amplify or sequence.

**Table A.5.** List of intrageneric genetic distance comparisons of our case-study samples with the kimura2 distance value for our long COI barcode.

| Species 1 | Species 2 | Kimura2 distance<br>in Long COI marker |
| --- | --- | --- |
| Araneidae Araneus angulatus<br>AM486 | Araneidae Araneus diadematus<br>DGMay02 | 0.139189801 |
| Cheiracanthiidae Cheiracanthium<br>punctorium TH318 | Cheiracanthiidae<br>Cheiracanthium virescens LG049 | 0.101477458 |
| Linyphiidae Tenuiphantes flavipes<br>TH333 | Linyphiidae Tenuiphantes<br>mengei LM288 | 0.09145557 |
| Linyphiidae Tenuiphantes flavipes<br>TH333 | Linyphiidae Tenuiphantes tenuis<br>DG027 | 0.097976615 |
| Linyphiidae Tenuiphantes mengei<br>LM288 | Linyphiidae Tenuiphantes tenuis<br>DG027 | 0.09256299 |
| Lycosidae Pardosa lugubris<br>DGMay14 | Lycosidae Pardosa hortensis<br>LM038 | 0.072967562 |
| Philodromidae Philodromus albidus<br>TH139 | Philodromidae Philodromus<br>cespitem LM444 | 0.096198916 |
| Philodromidae Philodromus albidus<br>TH139 | Philodromidae Philodromus<br>dispar AM659 | 0.102940409 |
| Philodromidae Philodromus albidus<br>TH139 | Philodromidae Philodromus<br>praedatus FM050 | 0.104003641 |
| Philodromidae Philodromus albidus<br>TH139 | Philodromidae Philodromus<br>aureolus MS1100 | 0.106342149 |
| Philodromidae Philodromus<br>aureolus MS1100 | Philodromidae Philodromus<br>praedatus FM050 | 0.052569794 |
| Philodromidae Philodromus<br>aureolus MS1100 | Philodromidae Philodromus<br>cespitem LM444 | 0.058680464 |
| Philodromidae Philodromus<br>aureolus MS1100 | Philodromidae Philodromus<br>dispar AM659 | 0.106219719 |
| Philodromidae Philodromus cespitum LM444 | Philodromidae Philodromus praedatus FM050 | 0.054354302 |
| Philodromidae Philodromus cespitum LM444 | Philodromidae Philodromus dispar AM659 | 0.101726261 |
| Philodromidae Philodromus dispar AM659 | Philodromidae Philodromus praedatus FM050 | 0.104110002 |
| Salticidae Evarcha arcuata LG017 | Salticidae Evarcha falcata AM278 | 0.092466694 |
| Theridiidae Enoplognatha ovata LG057 | Theridiidae Enoplognatha thoracica LM113 | 0.120842732 |
| Thomisidae Xysticus acerbus LG060 | Thomisidae Xysticus kochi FM138 | 0.095125415 |
| Thomisidae Xysticus acerbus LG060 | Thomisidae Xysticus lanio AM107 | 0.097398749 |
| Thomisidae Xysticus acerbus LG060 | Thomisidae Xysticus cristatus LM469 | 0.09849886 |
| Thomisidae Xysticus acerbus LG060 | Thomisidae Xysticus bifasciatus LK028 | 0.102892529 |
| Thomisidae Xysticus bifasciatus LK028 | Thomisidae Xysticus lanio AM107 | 0.101878734 |
| Thomisidae Xysticus bifasciatus LK028 | Thomisidae Xysticus kochi FM138 | 0.103790062 |
| Thomisidae Xysticus bifasciatus LK028 | Thomisidae Xysticus cristatus LM469 | 0.108312107 |
| Thomisidae Xysticus cristatus LM469 | Thomisidae Xysticus kochi FM138 | 0.036192062 |
| Thomisidae Xysticus cristatus LM469 | Thomisidae Xysticus lanio AM107 | 0.095344312 |
| Thomisidae Xysticus kochi FM138 | Thomisidae Xysticus lanio AM107 | 0.093093249 |

**Table A.6.**
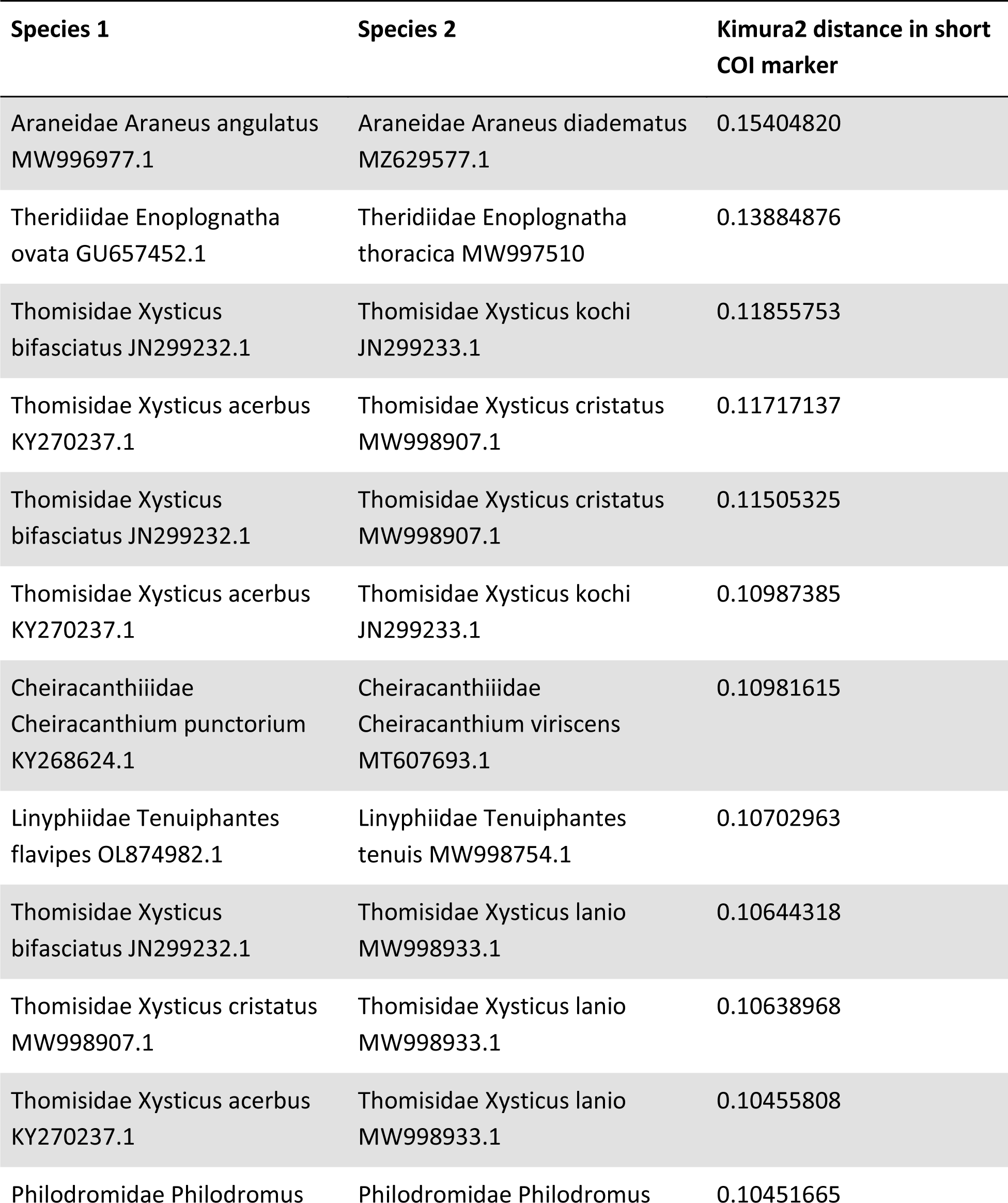
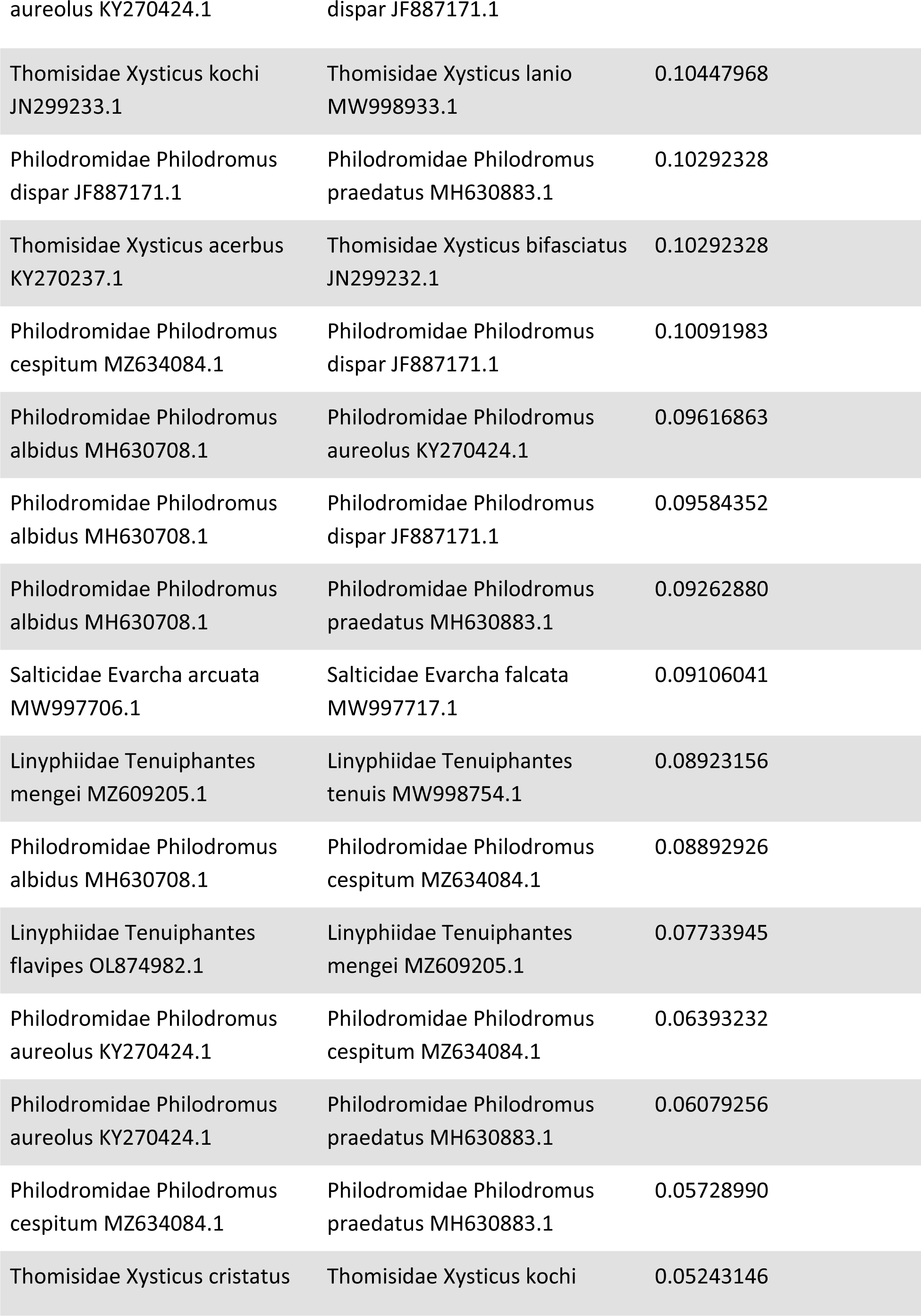
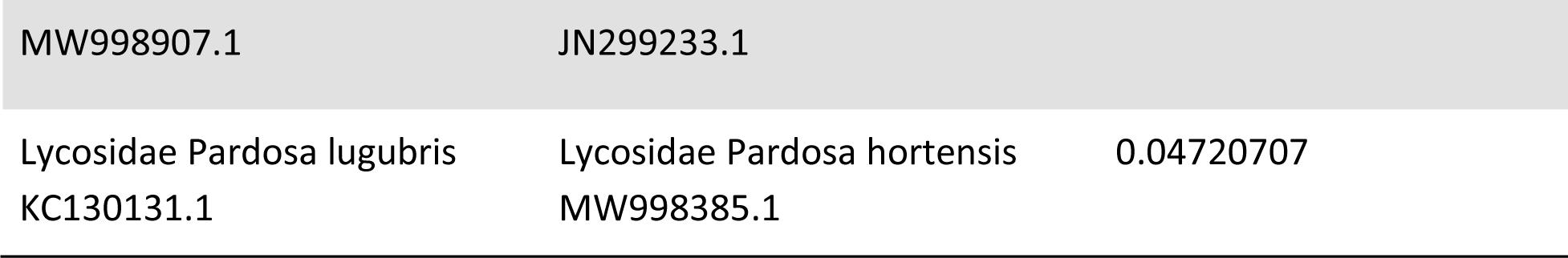
List of intrageneric genetic distance comparisons of our case-study samples with the kimura2 distance value for the short COI barcode downloaded from the NCBI database. The variable values next to the family, genus and species name of spiders are the accession numbers of sequences downloaded from the NCBI database.

**Figure A.1.**
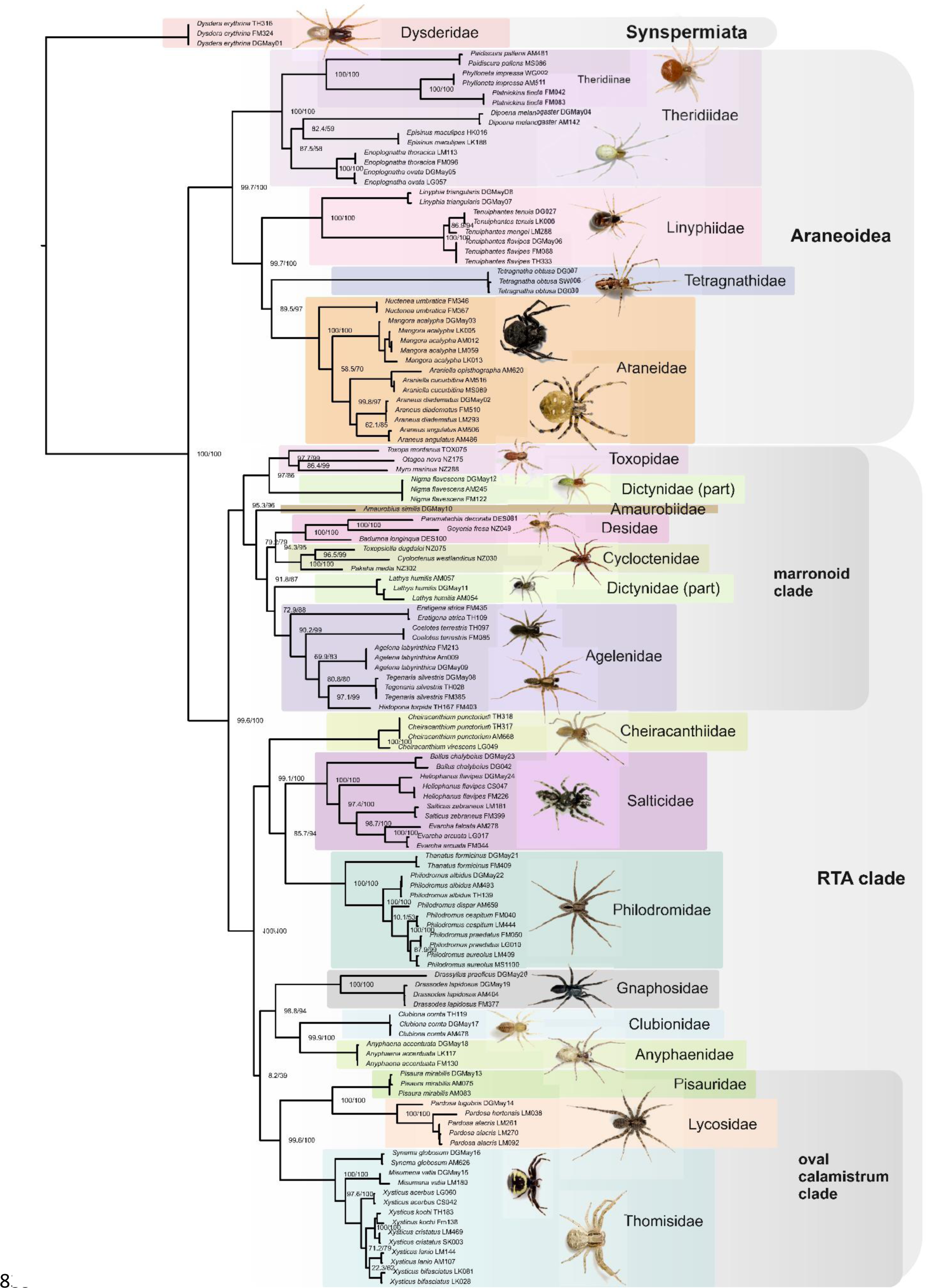
Maximum likelihood phylogeny inferred from the partitioned alignment of the three amplicons obtained using the newly designed primers COI_DGNS_F/R and CytB_DGNS_F/R, and the published primers 18S_F4 / 28S_R8 (Krehenwinkel et al. 2019), without Gblocks cleaning (for comparison with Fig. 2 where the sequences were Gblocks cleaned). Major clades widely recognized in spider systematics and recovered in previous multi-locus approaches are marked with color and labelled, though Dionycha is not recovered as monophyletic. Numbers at nodes give the ufb- and SH-aLRT node support values. The two specimens of *Histopona torpida* (Agelenidae) were merged into one sequence because one specimen only recovered the mitochondrial and the other only the nuclear amplicon.

**Figure A.2.**
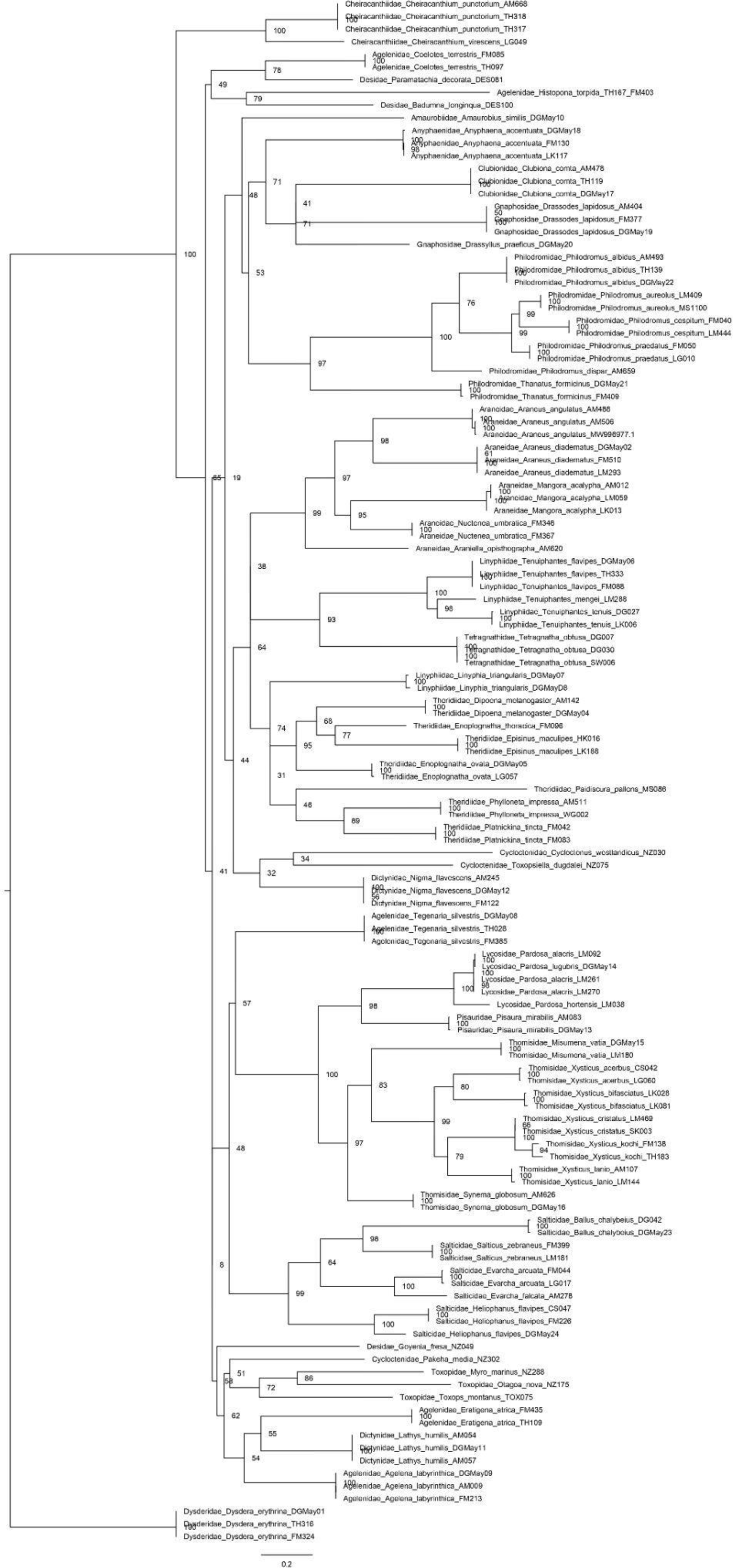
Maximum likelihood phylogeny inferred from the alignment of the 658-bp COI barcode. Amplicons obtained using the newly designed primers COI_DGNS_F/R were aligned with a downloaded barcode sequence of *Araneus angulatus* (Genbank accession number MW996977.1; Domènech et al. 2022), then trimmed to the 658-bp barcode region, after which the downloaded sequence was deleted from the alignment. The tree was generated in IQTree (Nguyen et al., 2015) with ultrafast bootstrapping (Minh et al., 2013) under the following partition model: codon 1 - HKY+F+R3; codon 2 - GTR+F+I+G4; codon 3 - TVM+F+I+G4, as determined by ModelFinder (Kalyaanamoorthy et al., 2017) in IQTree. Numbers at nodes give the ufb node support values. The topology, compared to the topology obtained from all three barcodes together (Fig. 2), is inferior with low support values. The topology does not support most of the clades widely recognized in spider systematics and recovered in previous multi-locus approaches.

**Figure A.3.**
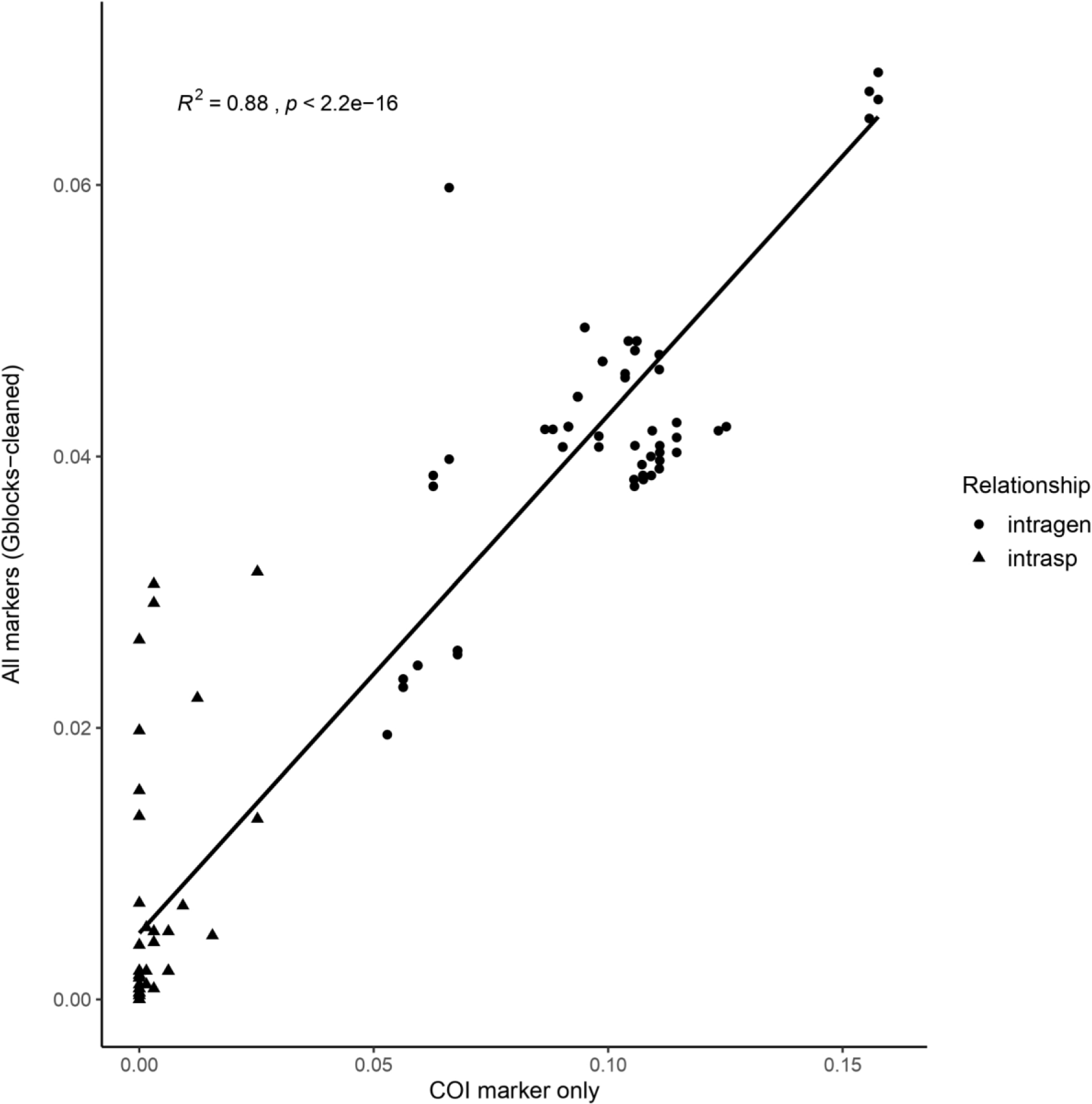
Relationship between intraspecific (triangles) and intrageneric (circles) genetic distances of the studies spider species. The *X*-axis shows the pairwise distance for our long COI amplicon, and the *Y*-axis depicts the corresponding pairwise distances for all of our three long-amplicon barcodes combined (COI, CytB and the 18S-28S region) and Gblocks- cleaned.

**Figure A.4.**
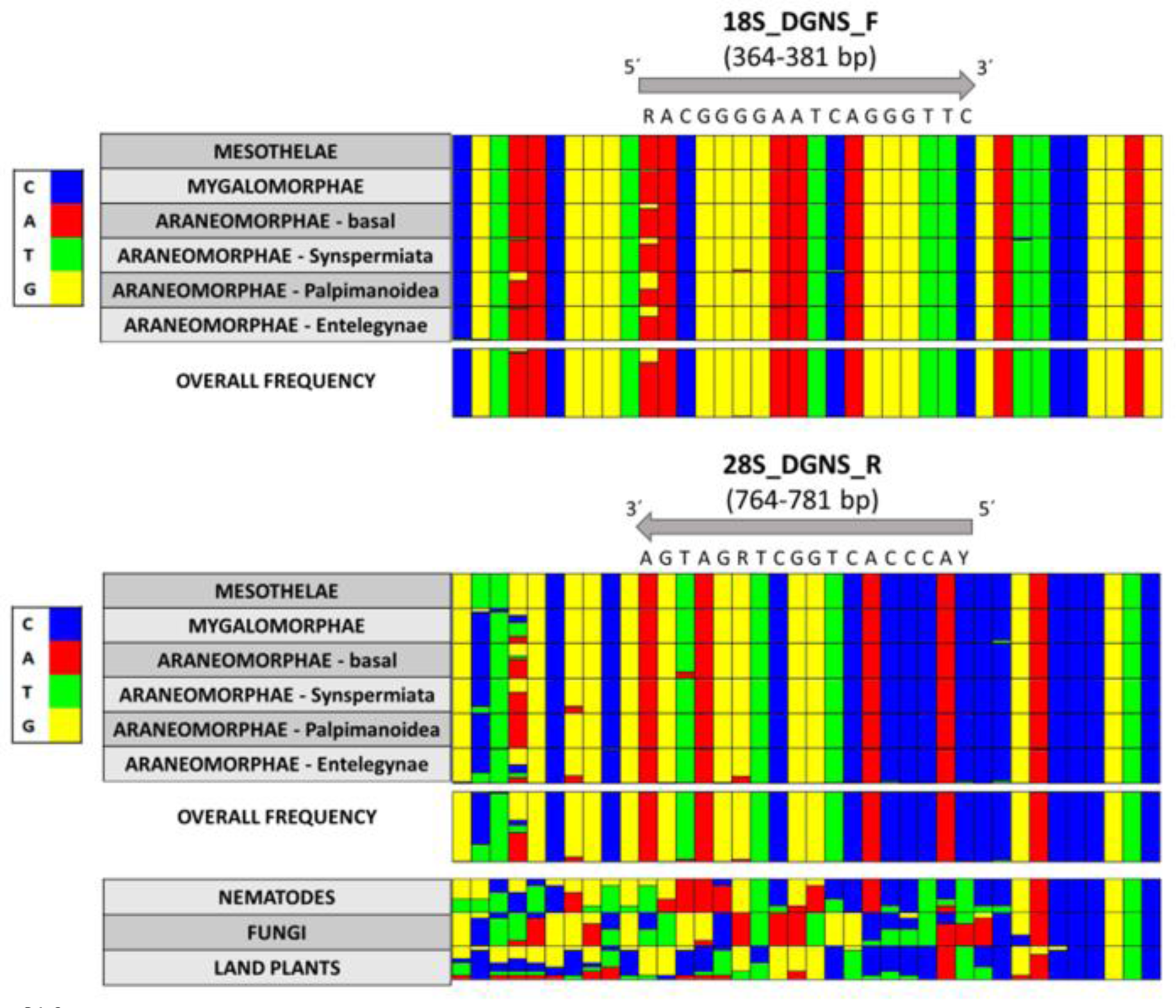
The nucleic acid composition of every base position inside the 18S_DGNS_F and 28S_DGNS_R primer region on the 18S and 28S gene (plus 10 nucleotides added to each side of the primer) from every major spider clade. The primer position in the gene is detailed below the primer name in brackets. The 18S and 28S gene sequences of a subsample of spider species were obtained from the NCBI database through the Wheeler et al. 2017 list. For better visibility, the sequences downloaded were merged into the major spider clades. The Araneomorphae - basal group contains species belonging to families Austrochilidae, Filistatidae, Gradungulidae and Hypochilidae. Here, we additionally provided the same region of 28S gene for several sequences downloaded for species belonging to the nematodes, fungi and land plants, to showcase how our primer is mismatching common contamination groups. The bases R and Y in the primer sequence represent the IUPAC nucleotide code for degenerate bases, which are common in universal primers. Note that the R primer is written in reverse complement, from 3’ to 5’. Figures were generated using the plot_alignments command in the R package PrimerMiner (Elbrecht & Leese 2017).

## Notes

### Competing Interest Statement

The authors have declared no competing interest.

