## Supplementary material for "Facilitating taxonomy and phylogenetics: An informative and cost-effective protocol integrating long amplicon PCRs and third generation sequencing": Script for sequencing data processing

#In Unix, unzip the fastq files and then concatenate them all into one:
gunzip *.fastq.gz
cat *.fastq > all_nanopore_files.fastq

#Demultiplex the files by index combo. Note, I'm making index edit distance (e) and primer edit distance (E) a bit smaller than recommended by 
#Pomerantz et al. 2022 to increase stringency:
python minibar.py dem.txt all_nanopore_files.fastq -F -e 2 -E 9


#Now use NGSpeciesID to simultaneously separate your samples by marker (inner primers) and generate consensus sequences:
#Example for a single sample for each marker:

NGSpeciesID --ont --consensus --q 10 --abundance_ratio 0.02 --m 3200 --s 800 --medaka --primer_file ribo_Kwink.txt --fastq sample_ALF_01B_Age_Ag_la_AM009.fastq --outfolder riboKwink_ALF_01B_Age_Ag_la_AM009
NGSpeciesID --ont --consensus --q 10 --abundance_ratio 0.001 --m 1050 --s 150 --medaka --primer_file COI.txt --fastq sample_ALF_01B_Age_Ag_la_AM009.fastq --outfolder COI_ALF_01B_Age_Ag_la_AM009
NGSpeciesID --ont --consensus --q 10 --abundance_ratio 0.001 --m 700 --s 150 --medaka --primer_file CytB.txt --fastq sample_ALF_01B_Age_Ag_la_AM009.fastq --outfolder CytB_ALF_01B_Age_Ag_la_AM009


#--q is quality threshold, --m is target amplicon length, --s is deviation from target length, --abundance_ratio is the minimum proportion
#of total reads a sequence must have in order to be considered for consensus formation.
#Note that the abundance ratio is lower for COI & CytB than for the ribo marker. I first tried all markers w/ abundance ratio of 0.02,
#but still had considerable dropout of COI & CytB (though ribo looked okay). Hence the change to 0.001 for COI & CytB. 

#Running NGSpeciesID individually for each specimen will take a lot of input, so do it in batch by running these scripts:
./consensus_ribo_abra.sh
./consensus_COI_abra2.sh
./consensus_CytB_abra2.sh

#Extract consensus sequences from folders and rename them by the parent folder and cluster ID:

#COI
ls */*fasta | awk '{print "cp "$0" "$0}' | sed 's/\//XXX/2' | sed 's/XXXconsensus_reference//' > cp_COI.sh
./cp_COI.sh

#Rename fasta headers by file name, and concatenate fasta files into one file for each marker:
awk '{if ($0~/>/) print ">"FILENAME;else print $0;}' COI*fasta > COI_All.fasta


#CytB
ls */*fasta | awk '{print "cp "$0" "$0}' | sed 's/\//XXX/2' | sed 's/XXXconsensus_reference//' > cp_CytB.sh
./cp_CytB.sh

awk '{if ($0~/>/) print ">"FILENAME;else print $0;}' CytB*fasta > CytB_All.fasta


#ribo
ls */*fasta | awk '{print "cp "$0" "$0}' | sed 's/\//XXX/2' | sed 's/XXXconsensus_reference//' > cp_ribo.sh
./cp_ribo.sh

awk '{if ($0~/>/) print ">"FILENAME;else print $0;}' ribo*fasta > ribo_All.fasta


#=========================================================================
##Download relevant sequences from NCBI (spider mitochondrial & ribosomal seqs, fungal 18S and 28S & mitochondrial). Then proceed:

#Make fasta files single-line:
awk '{if(NR==1) {print $0} else {if($0 ~ /^>/) {print "\n"$0} else {printf $0}}}' "18S_fungi".fasta > "18S_fungi_oneline".fasta
awk '{if(NR==1) {print $0} else {if($0 ~ /^>/) {print "\n"$0} else {printf $0}}}' "28S_fungi".fasta > "28S_fungi_oneline".fasta
awk '{if(NR==1) {print $0} else {if($0 ~ /^>/) {print "\n"$0} else {printf $0}}}' "mitochondrial_fungi".fasta > "mitochondrial_fungi_oneline".fasta
awk '{if(NR==1) {print $0} else {if($0 ~ /^>/) {print "\n"$0} else {printf $0}}}' "ribosomal_Araneae".fasta > "ribosomal_Araneae_oneline".fasta
awk '{if(NR==1) {print $0} else {if($0 ~ /^>/) {print "\n"$0} else {printf $0}}}' "mitochondrial_Araneae".fasta > "mitochondrial_Araneae_oneline".fasta

#Concatenate them together:
cat 18S_fungi_oneline.fasta 28S_fungi_oneline.fasta mitochondrial_fungi_oneline.fasta ribosomal_Araneae_oneline.fasta mitochondrial_Araneae_oneline.fasta > NCBI_seqs.fasta

#Change EOL encoding of big NCBI file to Windows-readable (skip this step if working on a Unix platform):
tr '\n' '\r' < NCBI_seqs.fasta > NCBI_seqs_Win.fasta

#Make this fasta file into a database:
makeblastdb -in NCBI_seqs_Win.fasta -dbtype nucl


#Search the file of all COI seqs against the database:
blastn -db NCBI_seqs_Win.fasta -query COI_All.fasta -outfmt "6 qseqid sseqid pident length mismatch gapopen qstart qend sstart send evalue bitscore staxids" -max_target_seqs 10 -out COI_blast.out


#Extract all the headers from the fasta files downloaded from NCBI. We'll match these up with the BLAST hits in the .out files:
awk 'sub(/^>/, "")' mitochondrial_Araneae_oneline.fasta > mitoAran_headers.txt
awk 'sub(/^>/, "")' ribosomal_Araneae_oneline.fasta > riboAran_headers.txt
awk 'sub(/^>/, "")' mitochondrial_fungi_oneline.fasta > mitofung_headers.txt
awk 'sub(/^>/, "")' 18S_fungi_oneline.fasta > 18Sfung_headers.txt
awk 'sub(/^>/, "")' 28S_fungi_oneline.fasta > 28Sfung_headers.txt

#Concatenate all the resultant files together so we end up with a list of taxa contained in our custom-made database:
cat *_headers.txt > NCBI_taxa.txt


#Now match up our BLAST results with the accession numbers (first column) of "NCBI_taxa.txt":
awk '{print "grep -E -m1 \""$2"\" NCBI_taxa.txt"}' COI_blast.out > grepCOI.sh
./grepCOI.sh > COI_blast_taxids.txt
paste COI_blast.out COI_blast_taxids.txt > COI_blast_taxa.txt

#Search the file of all CytB seqs against the database:
blastn -db NCBI_seqs_Win.fasta -query CytB_All.fasta -outfmt "6 qseqid sseqid pident length mismatch gapopen qstart qend sstart send evalue bitscore staxids" -max_target_seqs 10 -out CytB_blast.out
awk '{print "grep -E -m1 \""$2"\" NCBI_taxa.txt"}' CytB_blast.out > grepCytB.sh
./grepCytB.sh > CytB_blast_taxids.txt
paste CytB_blast.out CytB_blast_taxids.txt > CytB_blast_taxa.txt


#Search the file of all ribo seqs against the database:
blastn -db NCBI_seqs_Win.fasta -query ribo_All.fasta -outfmt "6 qseqid sseqid pident length mismatch gapopen qstart qend sstart send evalue bitscore staxids" -max_target_seqs 10 -out ribo_blast.out
awk '{print "grep -E -m1 \""$2"\" NCBI_taxa.txt"}' ribo_blast.out > grepribo.sh
./grepribo.sh > ribo_blast_taxids.txt
paste ribo_blast.out ribo_blast_taxids.txt > ribo_blast_taxa.txt


####Individually rerunning consensus on ALF_10G to recover COI:####
NGSpeciesID --ont --consensus --q 10 --abundance_ratio 0.0001 --m 1050 --s 250 --medaka --primer_file COI.txt --fastq sample_ALF_10G_The_Pa_pa_MS086.fastq --outfolder COI_ALF_10G_The_Pa_pa_MS086
#Extract and rename the fasta files:
ls */*fasta | awk '{print "cp "$0" "$0}' | sed 's/\//XXX/2' | sed 's/XXXconsensus_reference//' > cp_COI_ALF_10G.sh
#Rename the headers and concatenate the 4 resultant consensus seqs into one file:
awk '{if ($0~/>/) print ">"FILENAME;else print $0;}' COI*fasta > COI_ALF_10G.fasta 


####Demultiplexing & consensus formation for 25 test spiders:
python minibar.py dem.txt 30May_all.fastq -F -e 2 -E 9
./consensus_May.sh
ls */*fasta | awk '{print "cp "$0" "$0}' | sed 's/\//XXX/2' | sed 's/XXXconsensus_reference//' > cp_30May.sh
awk '{if ($0~/>/) print ">"FILENAME;else print $0;}' ribo*fasta > ribo_30May.fasta 
awk '{if ($0~/>/) print ">"FILENAME;else print $0;}' CytB*fasta > CytB_30May.fasta 
awk '{if ($0~/>/) print ">"FILENAME;else print $0;}' COI*fasta > COI_30May.fasta 
